# A story of nasal horns: A new species of *Iguana* Laurenti, 1768 (Squamata, Iguanidae) in Saint Lucia, St Vincent & the Grenadines, and Grenada (Southern Lesser Antilles) and its implications for the taxonomy of the genus *Iguana*

**DOI:** 10.1101/466128

**Authors:** Michel Breuil, Barbara Vuillaume, David Schikorski, Ulrike Krauss, Matthew N. Morton, Pius Haynes, Jennifer C. Daltry, Elisabeth Corry, Glenroy Gaymes, Joanne Gaymes, Nicolas Bech, Misel Jelic, Frédéric Grandjean

## Abstract

The Lesser Antilles, in the Eastern Caribbean, were long considered to have only two species in the genus *Iguana* Laurenti 1768: the Lesser Antillean iguana *Iguana delicatissima*, which is endemic to parts of the Lesser Antilles, and the common green iguana *Iguana iguana*, which also occurs throughout Central and South America. However, herpetologists and reptile collectors have pointed out strong physical differences between some of the island populations of *Iguana iguana* and those from the continent. Drawing on both morphological and genetic data, this paper describes a third species *Iguana insularis* sp. nov. from the southern Lesser Antilles, specifically the countries of Saint Lucia, St Vincent & the Grenadines, and Grenada. The new species is described based on the following unique combination of characters: Presence of high median and medium to small lateral horns on the snout; Small subtympanic plate not exceeding 20% of the eardrum size; Two or three scales of decreasing size anterior to the subtympanic plate; Fewer than ten small to medium triangular gular spikes; Medium sized dewlap; Low number of small to medium dispersed nuchal tubercles; Dark brown iris, with the white of the eye visible; Oval, prominent nostril; Short and relatively flat head; High dorsal spines; No swelling of the jowls in reproductively active males; Colour of head, body and tail changing from green to pale grey or creamy white in old adults; Vertical black stripes on body and tail, fading with age in some populations. This paper furthermore distinguishes two subspecies: *Iguana insularis insularis* from the Grenada Bank (comprising Grenada and the Grenadine islands), and *Iguana insularis sanctaluciae* from Saint Lucia. The form on the island of Saint Vincent has not been identified. Both subspecies are globally threatened by unsustainable hunting (including the pet trade) and by invasive alien species, including hybridization from invasive iguanas from South and Central America (*I. iguana* and *I. rhinolopha*, considered here as full species) that have become established in all three countries. The authors call for stronger measures to conserve the remaining purebred *Iguana insularis* sp. nov. throughout its range and for further research to identify other cryptic species and subspecies of *Iguana* in the Lesser Antilles.

## Introduction

The islands of the Lesser Antilles curve around the Eastern border of the Caribbean Sea between the Greater Antilles and South America and are noted for their rich diversity of endemic and globally threatened reptiles (Hedges 2018). Iguanas are among the most iconic animals of this archipelago, but opinions on their nomenclature and distribution have changed many times over the past few centuries.

Linnaeus (1758) described the common green iguana as *Lacerta iguana*, whereas Laurenti (1768) described the Lesser Antillean iguana as *Iguana delicatissima* and the common species as *I. tuberculata.* Both authors based their descriptions on drawings and specimens (Breuil 2002, 2013, 2016; Pasachnik *et al.* 2006). Later, Wiegmann (1834) described a third species with pronounced horns on its snout, *I. rhinolopha*, from Mexico. Duméril & Bibron (1837) subsequently recognised three *Iguana* species, using the names *I. tuberculata* for the common green iguana, *I. nudicollis* for the Lesser Antillean iguana and *I. rhinolopha* for the horned Mexican iguana, but found only two characters to separate *I. tuberculata* from *I. rhinolopha*.

The first mention of an iguana on the island of Saint Lucia was by Levacher (1834), but no information was given to determine the species. Breen (1844) commented that the iguanas were “an excellent sport for the native *chasseurs* (hunters)”. Bonnecour(t), a traveller in the mid-19^th^ Century, caught two specimens in Saint Lucia, which are now housed in the Museum National d’Histoire Naturelle (MNHN) in Paris, France (Breuil 2013, 2016). Duméril & Duméril (1851) recognised some morphological similarities between the horned specimens from Saint Lucia and the horned iguanas Duméril & Bibron (1837) had described in Mexico. This was corroborated by Boulenger (1885), who considered that a stuffed specimen from Saint Lucia belonged to the “variety” *rhinolopha*, together with specimens from Central America. Confusingly, however, Provancher (1890) reported the presence of *I. delicatissima* on Saint Lucia on the basis of a stuffed specimen observed in a house (Fig. 1), but his description and drawing of the specimen were too imprecise to confirm the species’identity. Dunn (1934) remarked that “The reports of *i. rhinolopha* from St Kitts and from Sta. Lucia is very strange… Possibly the horned mutation has appeared independently in that island”. Referring to *rhinolopha*, Barbour (1935) considered that: “The Antillean specimens are probably based on specimens incorrectly labelled as to locality” and added “If there really ever were iguanas on these islands, the mongoose has exterminated them”. (Small Asian mongooses, *Urva javanica*, were widely introduced to many of the Caribbean islands towards the end of the 19^th^ Century in an attempt to control rats and, possibly in Saint Lucia’s case, the venomous snake *Bothrops caribbaeus*: Des Vœux 1903, Nellis & Everard 1983).

**FIGURE 1.**
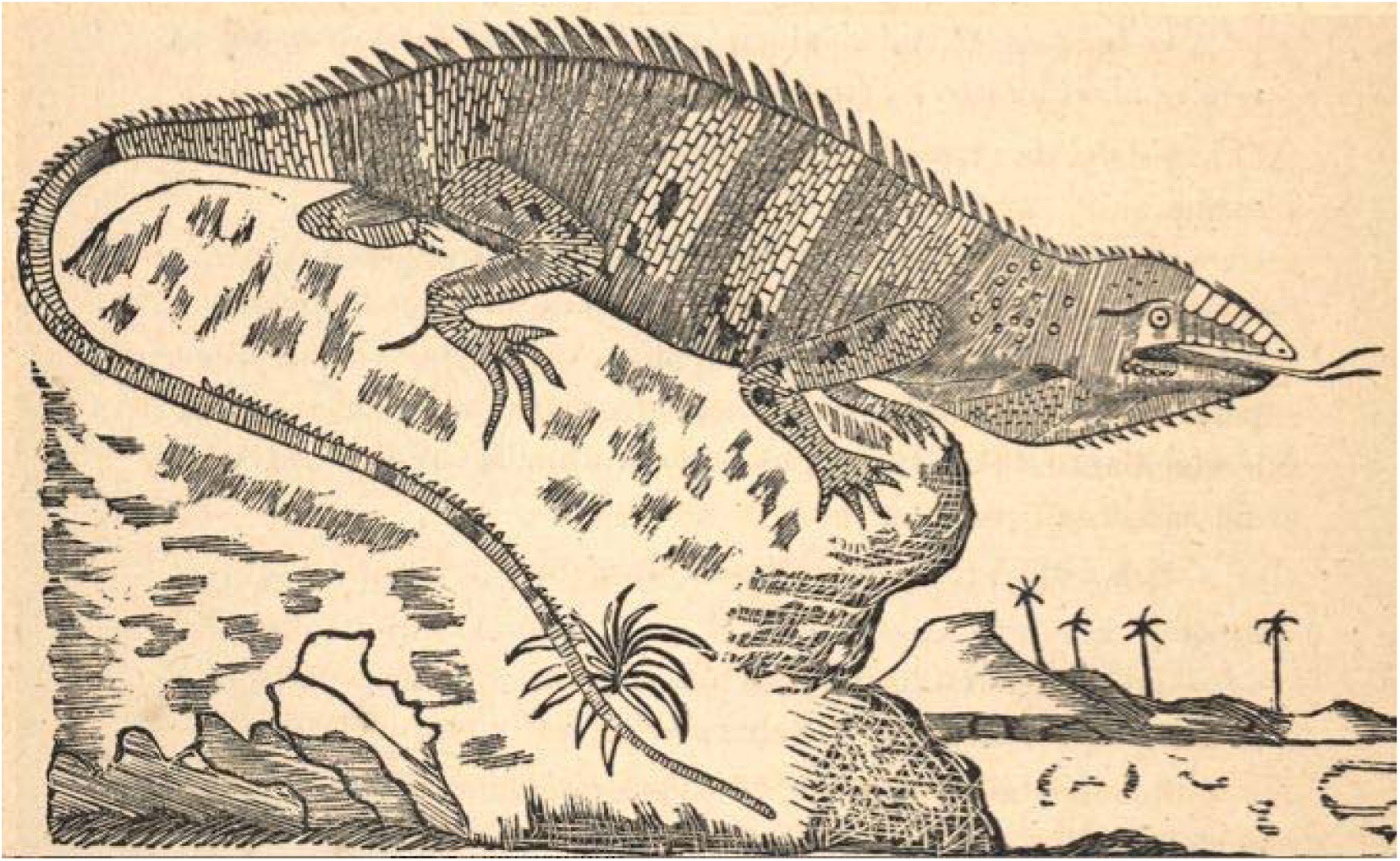
Drawing by Provancher (1890) of a stuffed iguana on Saint Lucia. Provancher identified it as *Iguana delicatissima* (see text), but the body and tail seem to have vertical black stripes, and there are small and scattered tubercular nape scales, and no subtympanic plate. There is no tympanum and no nasal horns on this drawing, which also shows a forked tongue

Farther south, Garmann (1887) remarked “the Grenada specimens are intermediate between *tuberculata* and *rhinolopha.* They have one prominent series of tubercles on the neck, and several scattered ones above the hind extremities. The tubercles on the snout are not so prominent as in *rhinolopha* from Central America, but the arrangement is the same. The tubercles on the neck are comparatively few as compared with those on Nicaraguan types”. Later, Barbour (1914) advised “a careful revision of these two species, made with the aid of extensive collections from many localities, will be necessary before their exact status can be settled”. He added: “That they are really distinct I have no doubt whatever, but as yet their ranges cannot be accurately defined. Stejneger suspects that if intermediates really do exist, they may be explained by the fact that the species have been carried by human agency ‘in innumerable instances’, and that the intermediates may be ‘hybrids from introduced stock, or because of their geographic distribution’ (*ex litt*.). I favour the latter explanation, as apparently the accidental introduction of vertebrates by human agency is a far rarer phenomenon than is often realized”. According to Dunn (1934) and Barbour (1935), Grenada was inhabited by *Iguana iguana iguana*.

Following Boulenger’s lead, Dunn (1934) recognised two species in the genus *Iguana*: *I. iguana* (with subspecies *iguana* and *rhinolopha*) and *I. delicatissima.* Lazell (1973) compared 139 “*I. iguana*” from Central and South America and the West Indies with 29 *I. delicatissima* collected between Martinique and Anguilla, and detected only one consistent difference between *I. iguana* and *I. delicatissima:* the existence in the former of a subtympanic plate at least 80% as large as the tympanum. Lazell (1973) rejected *rhinolopha* as a subspecies because he thought that the nasal horns were polytopic and polyphyletic characters, having observed them in iguanas from the Grenadines, Saint Lucia and parts of Central America. Based on their morphology, Lazell (1973) recognized three groups of *I. iguana* in the Lesser Antilles (Fig. 2): (1) The Northern group (Montserrat, Saba and, in the Greater Antilles, Saint Croix), distinguished by a higher proportion of melanistic individuals, large tubercular nape scales and dorsal crest spikes; (2) The Guadeloupe group (present only in Les Saintes and Basse Terre) which were “quite ordinary, and resemble those from northeastern South America. They may be patternless and/or grey: characteristics that are rare or absent in other parts of the range of *I. iguana*”; and (3) The Southern group between Saint Lucia and Grenada, characterised by vertical banding on the body and by nasal horns. Lazell considered the variation between the three groups to be clinal. For example, iguanas on the islands between Saint Lucia and Grenada have few tubercular nape scales; iguanas from Guadeloupe have more and larger ones; and those from the Northern group have very large and numerous tubercles.

**FIGURE 2.**
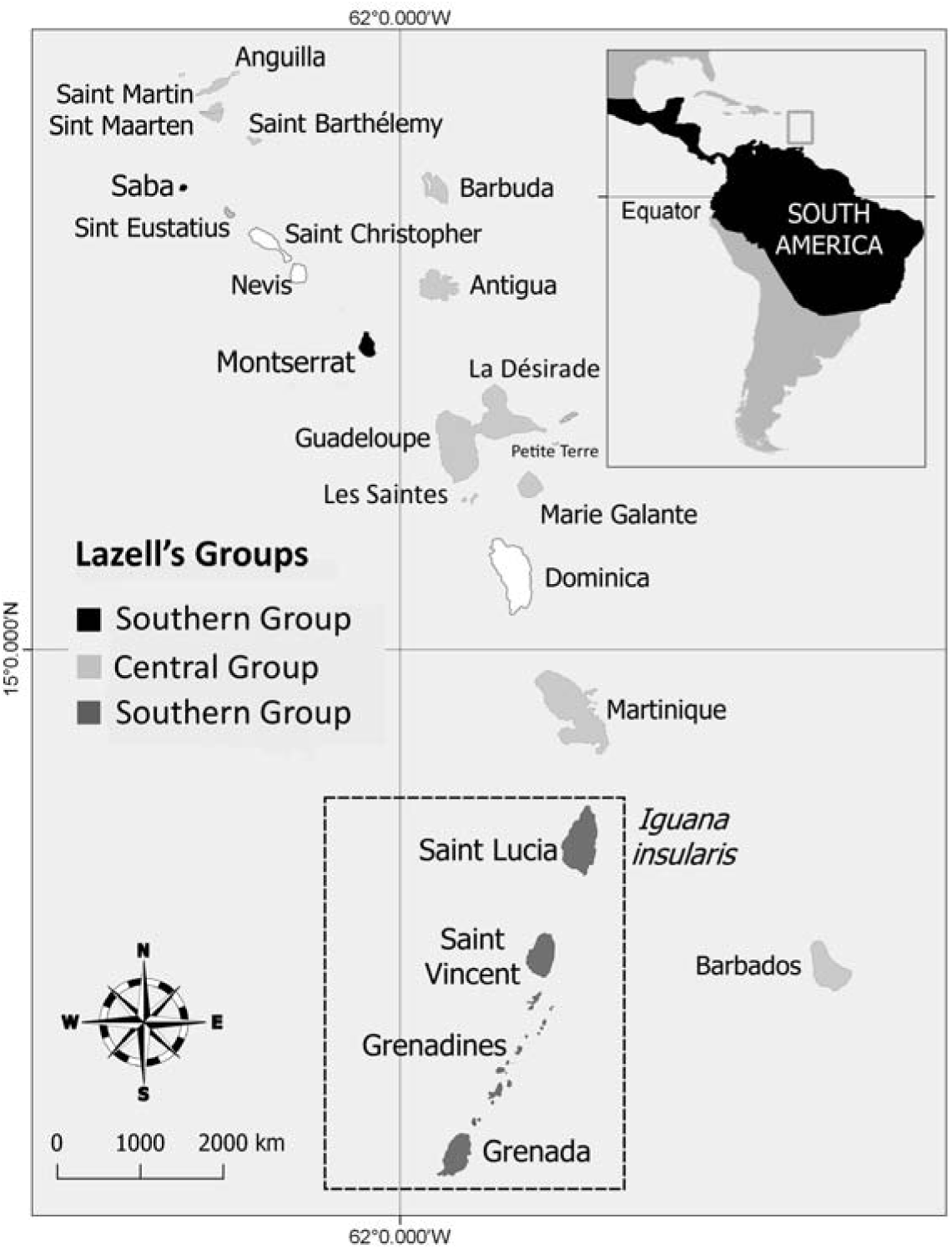
Distribution of the three iguana groups identified by Lazell (1973). Based on a clinal variation of their morphology, Lazell (1973) identified at the beginning of the Sixties 3 groups of the Common Iguana *Iguana iguana* that he thought to be native in Lesser Antilles. At that time, the central group was known to be present only in Guadeloupe (Les Saintes, Basse-Terre and Grande-Terre). This group is not native and is the descent of invasive common iguanas from French Guiana Breuil 2016; Vuillaume *et al.* 2015). Now, however, alien iguanas from Central and South America are present throughout most of this region (van den Burg *et al.* 2018) except Saint Christopher and Nevis, Dominica, Petite Terre and some satellites of Saint Barthélemy, Anguilla, and Martinique. The Southern Group is described in this work as a new species *Iguana insularis.*

Owing to their large size, striking colouration and horns, the Southern group is in demand for the international pet trade, and individuals are being smuggled out of these counties and marketed variously under the trade names “Saint Lucia iguana” and, from the Grenadines, “pink rhino iguana” and “white zebra rhino iguana” (Daltry pers. obs.; Noseworthy 2017).

In recent decades, this picture has been complicated by the discoveries both of hybridization between different species of *Iguana* and evidence of multiple invasions by iguanas from Latin America. When Lazell conducted his studies in the 1960s, both *I. delicatissima* and *I. iguana* were abundant and sympatric in Les Saintes (Guadeloupe), but not syntopic, and at that stage there was no evidence of one species displacing the other. It has since become clear that *I. iguana* is an invasive alien species in the Guadeloupean Archipelago (Breuil 2002, 2003), and that frequent interbreeding has taken place between *I. delicatissima* and *I. iguana* in Les Saintes, Grande-Terre and Basse-Terre, resulting in fertile and morphologically intermediate hybrids (Breuil 2002, 2003). Breuil (2013, 2016) reported that *I. iguana* was first introduced from French Guiana to Les Saintes in the mid-19^th^ Century, and Breuil (2009) and Breuil *et al.* (2010) explained how this species arrived on Basse-Terre. The unfortunate result of this introduction of *I. iguana* has been the elimination of the native *I. delicatissima* through hybridization and competition from Terre-de-Bas and Terre-de-Haut (Les Saintes) and Grande-Terre, a process that is continuing at present on Basse-Terre (Breuil *et al.* 2010; Vuillaume *et al.* 2015; Breuil 2013, 2016). More recently, *Iguana x Cyclura* hybrids have been recorded from Little Cayman Island (Moss *et al.* 2017), showing the lack of isolating mechanisms between these Caribbean genera.

From Guadeloupe, the alien iguanas have spread North and South. Father Pinchon brought *I. iguana* from Les Saintes to Martinique, and this invasive species now ranges over southern Martinique (Breuil 2011). Following Hurricane Luis in September 1995, dozens of iguanas were carried on rafts composed of logs, vegetation, house debris and garbage from Guadeloupe to Antigua, Barbuda and Anguilla (Daltry pers. obs.; Censky *et al.* 1998; Hodge *et al.* 2011): the Guadeloupe origin of these iguanas was inferred from their morphology (Breuil 1999). Both Anguilla and Barbuda now have growing populations of non-native *I. iguana* (Henderson & Breuil 2012). Additional iguanas have arrived in the West Indies as pets, putatively originating from breeding centres in Central America (Kraus 2009). These iguanas tend to be bigger than the Guadeloupean form, often with a yellow-orange iris, flat median horns on their snouts, big tubercular nape scales and a very big subtympanic plate. During the breeding season the males are often bright orange. These invasive alien iguanas have recently become established in the wild in Saint Martin, Saint Barthélemy, Martinique and Saint Lucia in the Lesser Antilles (Breuil 2013, 2016). Besides being transported as pets and by storms, invasive iguanas have also spread as stowaways. In 2018, for example, hybrid *iguana x delicatissima* were detected for the first time near the main port in Dominica, leading local conservationists to infer that *I. iguana* had recently arrived with shipping containers from other Caribbean islands (Jeanelle Brisbane, WildDominique, pers. comm.). In addition to having become the greatest threat to the Critically Endangered *I. delicatissima* in the Lesser Antilles (van den Burg *et al.* 2018), the invasive *I. iguana* have severe negative impacts for other wildlife and humans on other islands, *e.g.* Puerto Rico, the Caymans and Dominican Republic (Pasachnik *et al.* 2012; Falcón *et al.* 2012, 2013; M. Goetz, pers. comm.).

Invasive alien *I. iguana* were reportedly smuggled as juveniles into Saint Lucia in the late 1980s. Credible reports of free-living hatchlings in the vicinity of Soufriere (Southwest Saint Lucia) date back to 2007, putatively having escaped from a cage in the grounds of a hotel in Soufriere despite warnings from the Forestry Department to keep the animals and their offspring well secured. As the invasive iguanas began to multiply in spite of efforts by the Forestry Department and Durrell Wildlife Conservation Trust to catch and cull them, the native iguana population has become threatened by possible hybridization and competition (Morton & Krauss 2011; Krauss *et al.* 2014). At the time of writing, the invasive iguana population is growing in Southwest Saint Lucia, while the indigenous iguana population is less than 15 km away, in the Northeast. The Government of Saint Lucia recognises the indigenous Saint Lucia iguana as a distinct and fully protected species, despite it having long been regarded by the scientific community as merely a variant of *Iguana iguana*.

The status of the iguanas in Grenada and St Vincent & the Grenadines is less well understood because iguanas in both countries have long been regarded as a single, relatively abundant game species that can be hunted during the open season and freely transported by hunters and buyers within their respective borders. Specimens examined by the authors indicate that invasive *I. iguana* from Central and South America have invaded and multiplied on the larger islands at least, including the main islands of Grenada and St Vincent. Unaware of the possible diversity in iguana taxa, in 2005 the St Vincent & the Grenadines Forestry Department relocated 260 indigenous iguanas from Palm Island to the nearby Tobago Cays (also in the Grenadines) and the Kingstown botanical gardens on Saint Vincent in response to complaints from the owners of a resort on Palm Island that the iguanas were becoming a nuisance. During the hunting season (October to January inclusive), hunters commonly collect and transport live iguanas from the Grenadine islands to sell as bushmeat on Saint Vincent (G. Gaymes, pers. obs.).

As this narrative shows, understanding the distribution and taxonomy of iguanas in the Lesser Antilles has been repeatedly frustrated by differences of opinion among scientists on nomenclature and diagnostic characters, the accidental and deliberate movement of both native and invasive alien iguanas between islands, and hybridisation between members of the genus *Iguana.* There have, however, been some recent breakthroughs. Breuil (2002, 2013, 2016) identified more than 15 morphological characters to reliably differentiate *I. delicatissima* from *I. iguana* (see also Vuillaume *et al.* 2015). Breuil (2013, 2016) also proposed diagnostic characters to distinguish iguanas from Central America, South America, Montserrat, Saba and Saint Lucia. Malone & Davis (2004) and Stephen *et al.* (2013) provided preliminary genetic data that suggested that the Saint Lucia iguana forms an independent radiation in the Lesser Antilles, but they did not consider the horned iguanas on islands South of Saint Lucia, such as the Grenadines (which Lazell, 1973, had placed in the same phenotypic group as the Saint Lucia iguana). Vuillaume *et al.* (2015) studied the genetic variation of iguanas in the Lesser Antilles from Saint Lucia to Saint Martin (French West Indies). Following this work, Breuil *et al.* (in preparation) work on the genetic and morphological originality of the insular population of Saba and Montserrat.

The objectives of this paper are:

1. To clarify the taxonomic status of the iguanas of the Southern Lesser Antilles using new morphological and genetic data from Saint Lucia, St Vincent & the Grenadines, and Grenada.
2. To present new information on the distribution, threats and ecology of this group, and recommendations for their conservation.

## Materials and methods

Morphological, molecular (*i.e.* mitochondrial DNA and microsatellites markers), and biological data were used to characterise the iguanas of Saint Lucia, Grenada and St Vincent & the Grenadines, and compare them to other populations of *Iguana iguana sensu lato*.

### Morphological analysis

The morphological characters used to examine the iguanas followed Breuil (2013, 2016), most of which are meristic characters that were easy to record from digital pictures taken by the authors of wild individuals and from specimens at the Museum of Comparative Zoology (MCZ) in Harvard, USA, and the Muséum National d’Histoire Naturelle (MNHNP) in Paris, France.

We also examined photographs of *Iguana* found on the Internet using the Google Images search engine for the islands of Saint Vincent, the Grenadines and Grenada. The use of Internet images for taxonomic research was advocated by Leighton *et al.* (2016) for studying spatial patterns in phenotypic traits that are objective, binary and easy to see, irrespective of the angle, to supplement fieldwork. To identify diagnostic characters, we retained only pictures that reported precise localities, and eliminated areas known to have iguanas introduced from Central and South America.

### Molecular analysis

#### Collection and preparation of genetic material

Genomic DNA was isolated from 39 specimens from tissue, shed skin and/or blood samples, using the QIAamp DNA Mini Kit (QIAGEN, Deutschland) and following the manufacturer’s recommendations (Table 1). Not all specimens were used for both mtDNA and microsatellites analysis. Samples from the Lesser Antilles were collected by the authors and from French Guiana by François Catzefis (CNRS, France) and Benoît de Thoisy (Institut Pasteur, Cayenne, French Guiana).

**Table 1.**
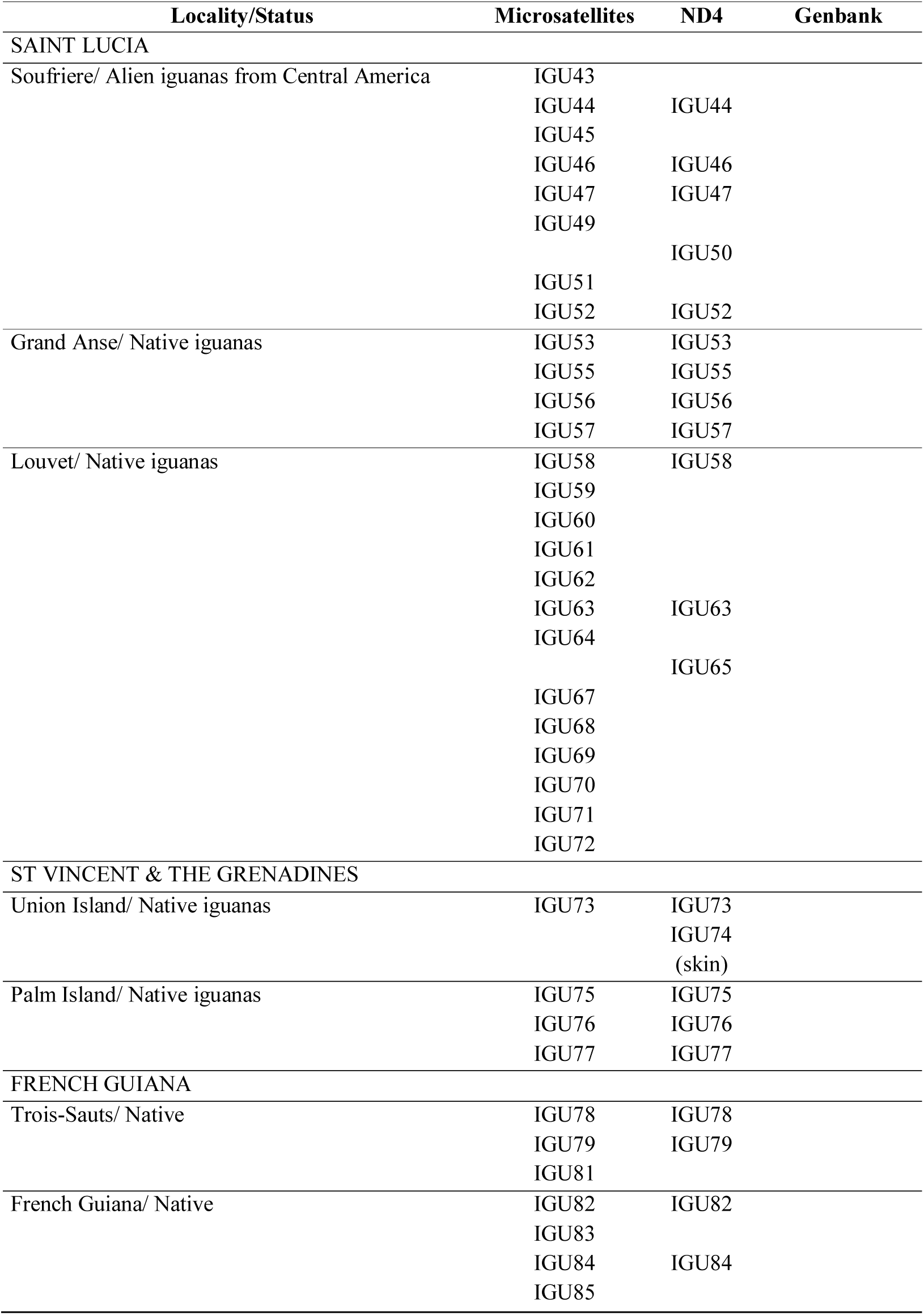
Genetic samples

#### Mitochondrial DNA (ND4)

903 base pairs (bp) of the ND4 mitochondrial DNA gene were amplified using primer pair 5’-CAC CTA TGA CTA CCA AAA GCT CAT GTA GAA GC-3’ and 5’-GCT TCT ACA TGA GCT TTT GGT AGT CAT AG-3’. A Qiagen multiplex PCR kit was used to conduct each PCR, with a total reaction volume of 25 μL containing 20ng DNA template, 12.5 μL Qiagen PCR Master Mix, 2.5 μL Qiagen Q-solution, and 2.5 μL primer mix at 10μM each. PCR reactions were carried out in a SimpliAmp thermal cycler under the following conditions: initial denaturation at 95°C for 15 min, followed by 35 cycles of denaturation at 95°C for 30 s, annealing at 52°C for 30 s, and extension at 72°C for 90 s, with a final extension step at 60°C for 30 min. The amplification was verified by electrophoresis using LabChip GX Analyser (Caliper Life Sciences, USA) and successful PCR products were then vacuum-purified using MANU 30 PCR plates (Millipore) before being sequenced using the ABI Big-Dye Terminator v3.1 Cycle Sequencing Kit (Thermo). Cycle sequencing reactions were finally purified with Sephadex G-50 Fine (GE Healthcare) and sequenced on an ABI 3130xl DNA sequencer (Applied Biosystems). Sequence chromatograms were analyzed in SEQUENCHER (v5.3; Gene Codes Corp., Ann Arbor). Sequence alignment was prepared with MAFFT (v7.187; Katoh *et al.* 2005). For comparisons we chose a 340-bp fragment of the ND4 locus common to all our specimens.

#### Microsatellites

This data set comprised 36 individuals representing seven insular and continental populations (Table 1). A panel of 17 microsatellite markers were amplified as described by Valette *et al.* (2012) and Vuillaume *et al.* (2015).

#### Phylogenetic analysis

For ND4 analysis, we aligned our ND4 sequences from the 21 specimens sampled by the authors (Table 1) with Genbank sequences obtained from previous studies (Malone *et al.* 2000; Malone & Davis 2004; Stephen *et al.* 2013; Martin *et al.* 2015). Phylogenetic trees were constructed using the Maximum Likelihood (ML) and Bayesian Inference (BI): The best-fit evolutionary model was calculated using the Bayesian Information Criterion in jModelTest2 (version 2.1.6; Darriba *et al.* 2012) while the ML analysis was conducted in MEGA6 (Tamura *et al.* 2013) based on the best-model obtained in jModeltest. Initial tree(s) for the heuristic search were obtained by applying the Neighbour-Joining method to a matrix of pairwise distances estimated using the Maximum Composite Likelihood (MCL) approach. A discrete Gamma distribution was used to model evolutionary rate differences among sites (5 categories). The BI was performed using MrBayes 3.2 (Ronquist *et al.* 2012) on the Cipres Science Gateway. Two independent runs with four MCMC chains were carried out for 50 million generations. The temperature parameter was set to 0.2 and chains were sampled every 5,000 generations. The first 12.5 million generations were discarded as burn-in. The effective sample sizes of parameters were checked using TRACER 1.5 (Drummond & Rambaut 2007) and the convergence of runs was checked using AWTY (Nylander *et al.* 2008). Supported nodes in phylogram were indicated with bootstrap values P ≥ 70 in ML and posterior probabilities (pp) values ≥ 0.95 in BI.

The Median-Joining (MJ) haplotype network (Bandelt *et al.* 1999) was constructed to analyze inter- and intraspecific relations among *Iguana* lineages. The MJ network was calculated and drawn using PopART (Population Analysis with Reticulate Trees) v1.7 (Leigh & Bryant 2015).

#### Genetic diversity

We tested departures from Hardy-Weinberg expectations and linkage disequilibria using exact tests based on the Markov chain (1,000 permutations) with the software FSTAT v. 2.9.3.2 (Goudet 2001). We adjusted the levels of significances for multiple tests using the standard Bonferroni correction (Rice 1989). We assessed the polymorphism over all loci for each population, computing allelic richness (AR), expected heterozygosity (He), allelic frequencies and inbreeding coefficient (Fis) (Weir & Cockerham 1984) using FSTAT v. 2.9.3.2 (Goudet 2001) with 1,200 permutations. The allelic frequencies allowed us to deduce private alleles for each population.

#### Genetic structure

We estimated pairwise fixation index (F_ST_) values between populations (Weir & Cockerham 1984) using FSTAT v. 2.9.3.2 (Goudet 2001). Their associated significance was computed and tested using global tests implemented in FSTAT v. 2.9.3.2 (Goudet 2001) with a level of significance adjusted for multiple tests using the standard Bonferroni correction. In addition, relationships among populations were evaluated with a Factorial Correspondence Analysis (FCA) based on individual genotypes and using the FCA procedure implemented in GENETIX v. 4.05.2 (Belkhir *et al.* 2004). We also accessed the genetic structure using the individual-based approach implemented by the software STRUCTURE (Pritchard *et al.* 2000). This Bayesian clustering approach estimated both the number (K) of genetic cluster(s) and the admixture coefficient of individuals to be assigned to the inferred clusters. We choose the admixture model and the option of correlated allele frequencies among populations. As recommended by Evanno *et al.* (2005), we replicated 20 independent runs for each value of K (with K varying from 1 to 10) with a total of 1 million iterations and a burn-in of 10,000. To determine the number of genetic clusters from STRUCTURE analyses, we used the STRUCTURE HARVESTER program (Earl & VonHoldt 2011) to compare the mean likelihood and variance per K values computed from the 20 independent runs.

## Systematic analysis

Based on morphological and genetic analysis, the Southern group of iguanas first identified by Lazell (1973) is herein recognized as a new species endemic to Saint Lucia, St Vincent & the Grenadines, and Grenada, with at least two subspecies.

### *Iguana insularis* new species

Southern Antillean horned iguana Figs 3-6.

**FIGURE 3.**
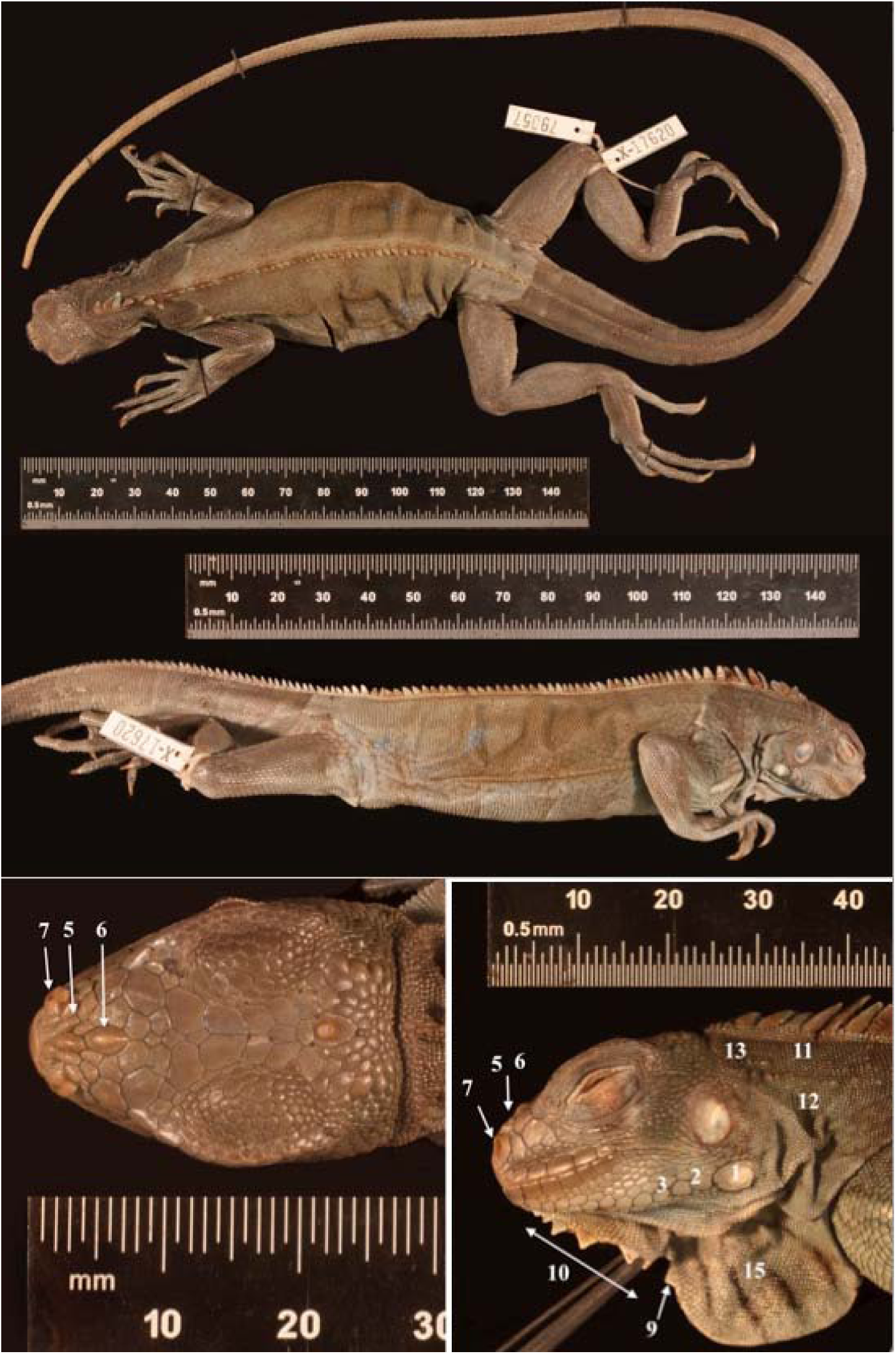
Holotype of *Iguana insularis insularis* ssp. nov. MCZ X-17620/R-79057 © Museum of Comparative Zoology, Harvard University. Specimen in alcohol with discolouration. Annotations: 1. Small size of subtympanic plate ± 10-20% of the eardrum. 2. Two to three scales of decreasing size anterior to subtympanic plate. 3. Juxtaposed elongated sublabial scales 5. Median and lateral horns on the snout. 6. Horns with enlarged bases. 7. Oval and prominent nostrils. 9. Flat and triangular gular spikes. 10. Six gular spikes. 11. Scattered nuchal tubercles. 12. Low number of nuchal tubercles. 13. Small size of nuchal tubercles. 15. Dewlap of medium size.

**FIGURE 4.**
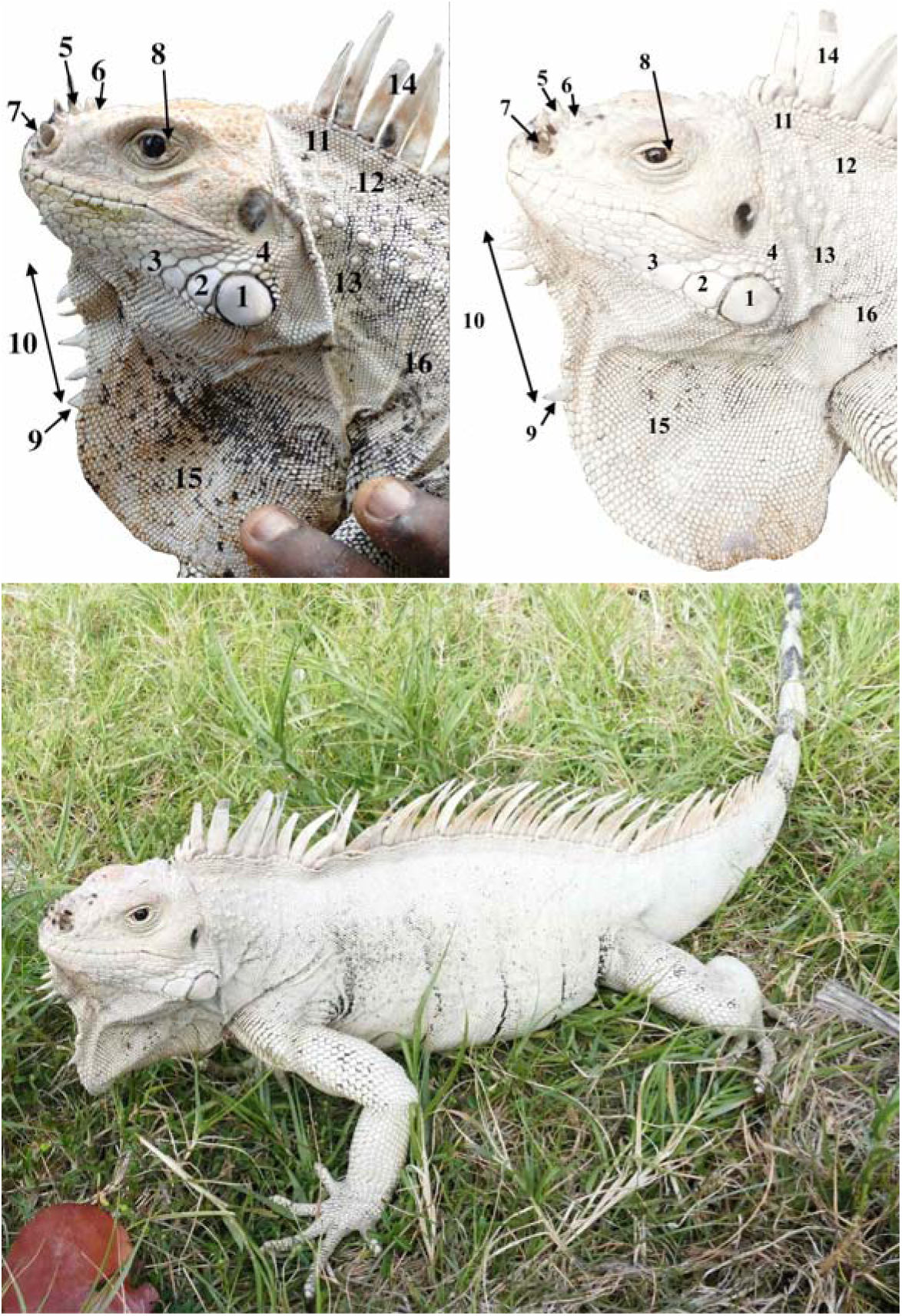
Examples of adult male *Iguana insularis insularis* ssp. nov. Top left. Adult male sequenced under the number IGU75 (SVL 41 cm); Top right and bottom, adult male sequenced under the number IGU77 (SVL 45 cm). Annotations: 1. Small size of subtympanic plate ± 10-20% of the eardrum. 2. Two or three scales of decreasing size anterior to subtympanic plate. 3. Juxtaposed elongated sublabial scales. 4. No apparent swelling of the jowls in breeding males. 5. Median and lateral horns on the snout. 6. Horns with enlarged bases. 7. Oval and prominent nostrils. 8. Brown eyes with visible white. 9. Flat and triangular gular spikes. 10. Seven or eight gular spikes. 11. Scattered nuchal tubercles. 12. Low number of nuchal tubercles. 13. Small size of nuchal tubercles. 14. Orange in dorsal scales in breeding animals. 15. Creamy white dewlap of medium size. 16. Creamy white body with faint to no black banding in old individuals.

**FIGURE 5.**
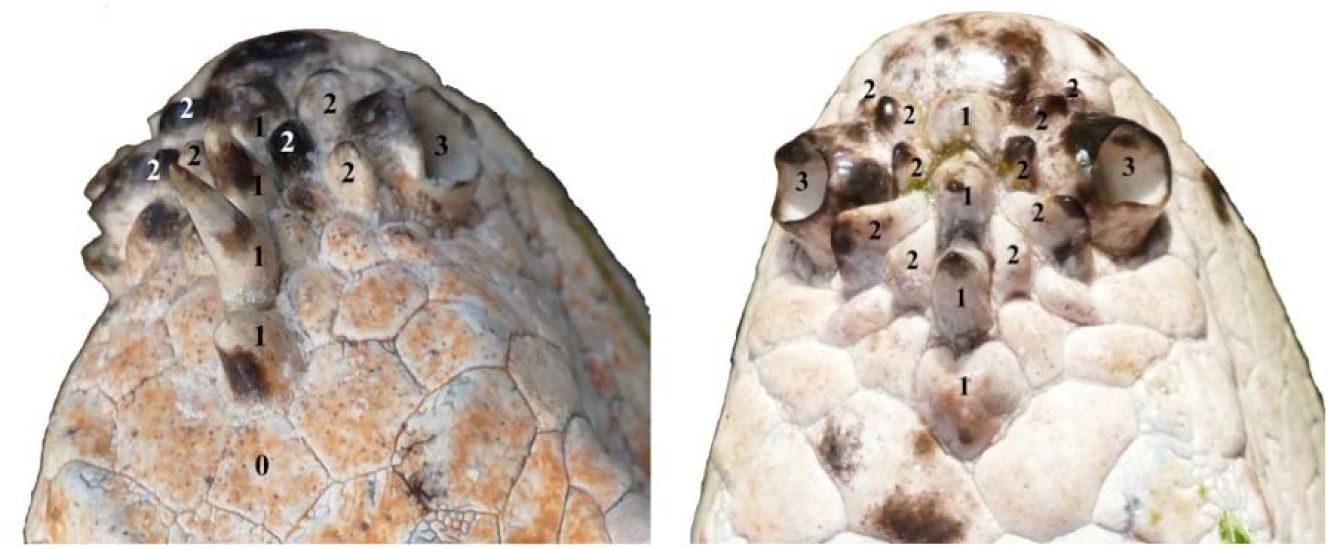
Nasal horns of *Iguana insularis insularis* ssp. nov. View of the snout of IGU75 (left) and IGU77 (right) from Palm Island (same individuals as Fig. 4). Annotations: 0. Frontal scale not developed into a horn. 1. Median horns with enlarged bases. 2. Lateral horns. 3. Oval prominent nostril. Note the differing forms and disposition of cephalic scales, and that IGU75 (a younger male) has flat scales whereas IGU77 (an older, larger male) has more prominent scales.

**FIGURE 6.**
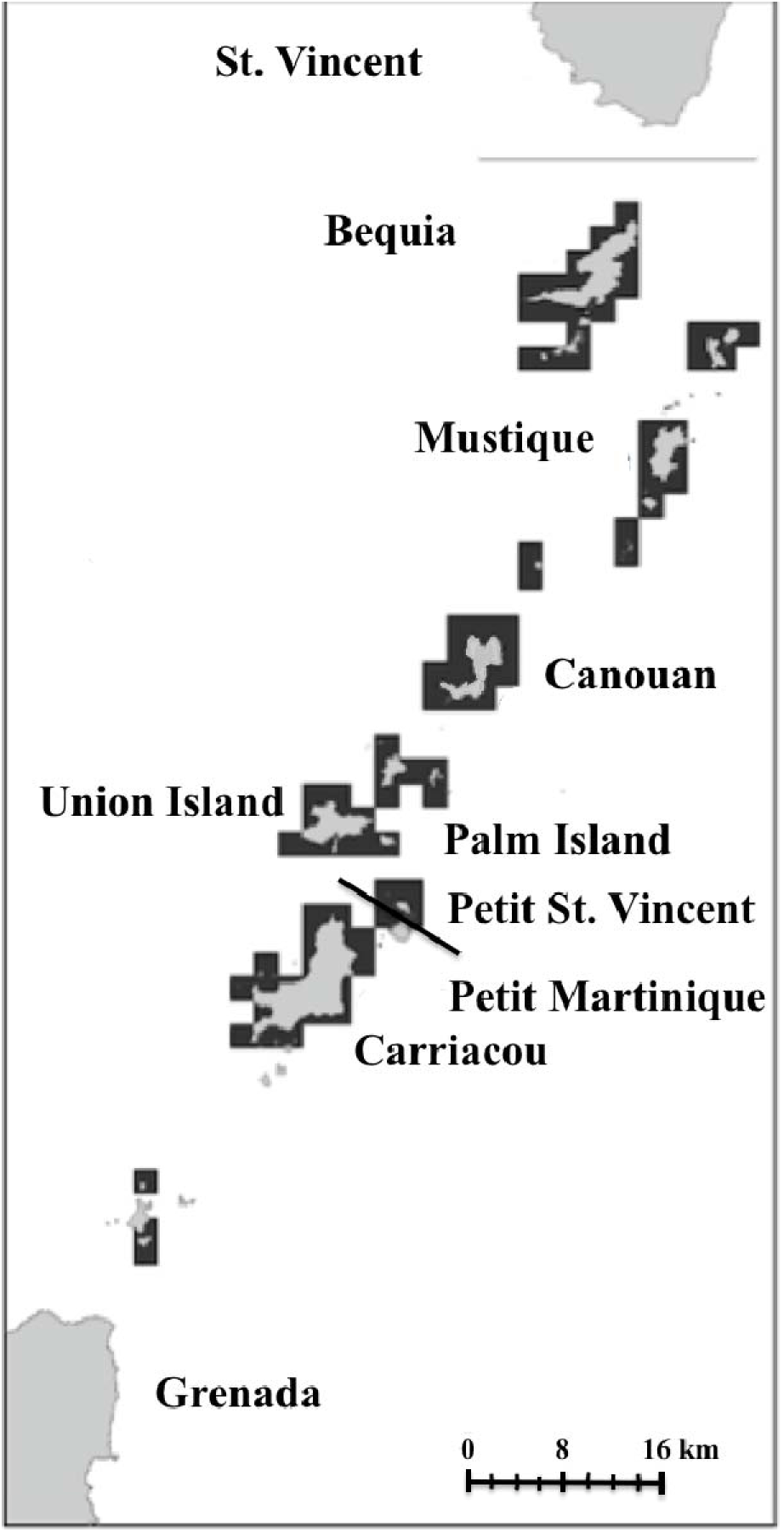
Distribution of iguanas in the Grenadine islands. Locations are mapped to the nearest 2×2 km square representing groups of islands in the Grenadines. We have deliberately avoided being specific to protect the animals (see Auliya *et al.* 2016). Note that there are alien iguanas on some islands and not all of the island clusters shown here have purebred populations of *Iguana insularis insularis.* We have no confirmed specific localities for the main island of Grenada, although a museum specimen (MCZ R-79747) confirms this subspecies occurred here. Henderson & Breuil (2012), Henderson & Powell (2018), Baldwin (2012) and Baldwin & Mahon (2011), G. Gaymes and J. Daltry (pers. obs.; the Grenadines). While iguanas are present on St. Vincent, these have not been identified to subspecies level and cannot be assumed to be identical to those on the Grenada Bank. The grey line just south of St Vincent marks the brake between the St. Vincent Bank to the north and the Grenada Bank to the south. The black line between Petit Saint Vincent and Petit(e) Martinique shows the political boundary between St. Vincent and the Grenadines to the north and Grenada to the south. The Grenadine islands form an archipelago from the south of St. Vincent to the north of Grenada.

This new species is characterized by the following combination of features found in both subspecies:

– Presence of median and lateral horns on the snout that are generally enlarged at the base;
– Small subtympanic plate, not exceeding 20 % of the high of the tympanum;
– Two or three scales of decreasing size anterior of the subtympanic plate;
– Not more than 8 small to medium triangular gular spikes, exceptionally 10;
– Dewlap of medium size;
– Low number of small to medium dispersed nuchal tubercles;
– Dark brown iris (never yellow to orange), with the white of the eye visible except in the juveniles;
– Oval prominent nostril, sometimes triangular;
– Short and relatively flat head;
– High dorsal spines;
– No swelling of the jowls in reproductively active males.
– Body of juveniles and young adults is predominantly bright green with 6-8 black vertical bands; The body becomes very pale (almost white or cream white) in old individuals and the vertical bands either fade (subspecies *insularis*) or remain black (subspecies *sanctaluciae*).
– Black bands on the tail, which typically remain conspicuous throughout life.

The new species has at least two subspecies, both described below: (1) The nominate subspecies on the Grenada Bank (comprising the main island of Grenada and the islands of the Grenadines), and (2) A subspecies restricted to the island of Saint Lucia. The most conspicuous difference between the two subspecies is the colouration of older adults. In both species the body colouration becomes pale, almost white, with age. However, in the nominate subspecies, the dark banding on the body is thinner and fades with age, whereas in Saint Lucia population, the bands are thicker and remain black. In both subspecies, the dewlap in the juveniles is green but in the Saint Lucia population the dewlap becomes entirely black with age whereas in the Grenada Bank iguanas the dewlap becomes cream white. These morphological differences prompted us to describe two subspecies on the Grenada Bank and Saint Lucia respectively, as presented below. Note that the form of *I. insularis* on St Vincent remains unverified due to lack of known pure-bred specimens from here.

### *Iguana insularis insularis* new subspecies

Grenadines horned iguana, pink rhino iguana

#### Holotype

The holotype of *Iguana insularis insularis* housed in MCZ under the numbers X-17620/R-79057 (Fig. 3). This specimen was caught by James Lazell on Bequia, St Vincent & the Grenadines (10 April 1964).

***Sex*:** Undetermined.

***Age*:** Young, possibly 2 years old, based on its size.

***Morphological measurements*:** Total length: 51.5 cm, SVL: 13.5 cm, tail length: 38 cm.

Height and width of the left subtympanic plate: 3.2 mm, 4.3 mm.

***Meristics*:** Number of gular spikes, 5 medium + 2 small. Number of dorsal spikes to cloaca: 54 ± 1.

#### Paratypes

Two other young specimens MCZ X-17619/R-79056 and X-17621/R-79058) from the same location and the same collector.

#### Diagnosis of *Iguana insularis insularis* (Figs 4, 5)

We define the typical morphology of this new taxon based on our own observations on both adults and juvenile specimens on Palm and Union Islands (St Vincent & the Grenadines), complemented by the specimens from MCZ R-79056-57-58 collected on Bequia (also in St Vincent & the Grenadines) and R-79747 from Sandy Bay, Grenada. The latter four are young individuals with SVL from 128 mm to 135 mm, and thus lack some details specific to adults.

The iguanas from the Grenada Bank, including the Grenadines, are characterised by the following association of characters in adults compared with iguanas from Saint Lucia (*I. insularis sanctaluciae* ssp. nov.).

– In most old adults (both males and females), the green colouration and black bands fade to an almost uniform light cream to nearly white, except on the posterior end of the tail where the black banding persists;
– In old adults, the head is nearly light cream to white;
– The dewlap is predominantly white but may have some black scales;
– There are no black margins on the subtympanic plate and on the sublabial scales;
– The snout has 2 to 5 median horns (usually 3 or 4) and 2 to 6 less developed lateral horns on each side;
– The horns may or may not remain black throughout the animal’s life;
– There are light yellow scales on the head and on the dewlap in old adults;
– The tips of the dorsal spikes of mature adults during the breeding season are light yellow to light orange;
– The anterior part of the dewlap is rounded.

#### Size

The largest purebred *I. insularis insularis* measured by the authors had an SVL of 45 cm (IGU77, an adult male on Palm Island, Fig. 4). Its tail was incomplete.

Another large individual fitting the morphology of this subspecies (but not yet genetically tested) on Petit Bateau had a total length of 136 cm.

#### Geographical distribution (Fig. 6)

Of the c. 30 islands of Grenada Bank, including the Grenadine islands and the main island of Grenada, 26 have been reported to have iguanas (Henderson & Powell 2018). The entire bank is inferred to have been originally inhabited by *I. insularis insularis* but morphological and genetic data indicate that several islands, including the main island of Grenada, have had incursions of *I. iguana* from South American and/or Central American lineages.

From our collection of photographs of specimens captured by the authors and obtained from internet searches, it is clear that most Grenadine islands still have the indigenous white, horned and more or less black-banded phenotype, but there is the Central America phenotype with various hybrids among them that make it difficult to confirm which islands still have purebred populations of this subspecies. Further genetic testing is required to accurately map the present distribution of *I. insularis insularis*.

#### Etymology

The specific name and thus the subspecific name refer to the numerous islands in the Southern Antilles where the new species lives.

### *Iguana insularis sanctaluciae* new subspecies

Saint Lucia horned iguana Figs 7-11.

**FIGURE 7.**
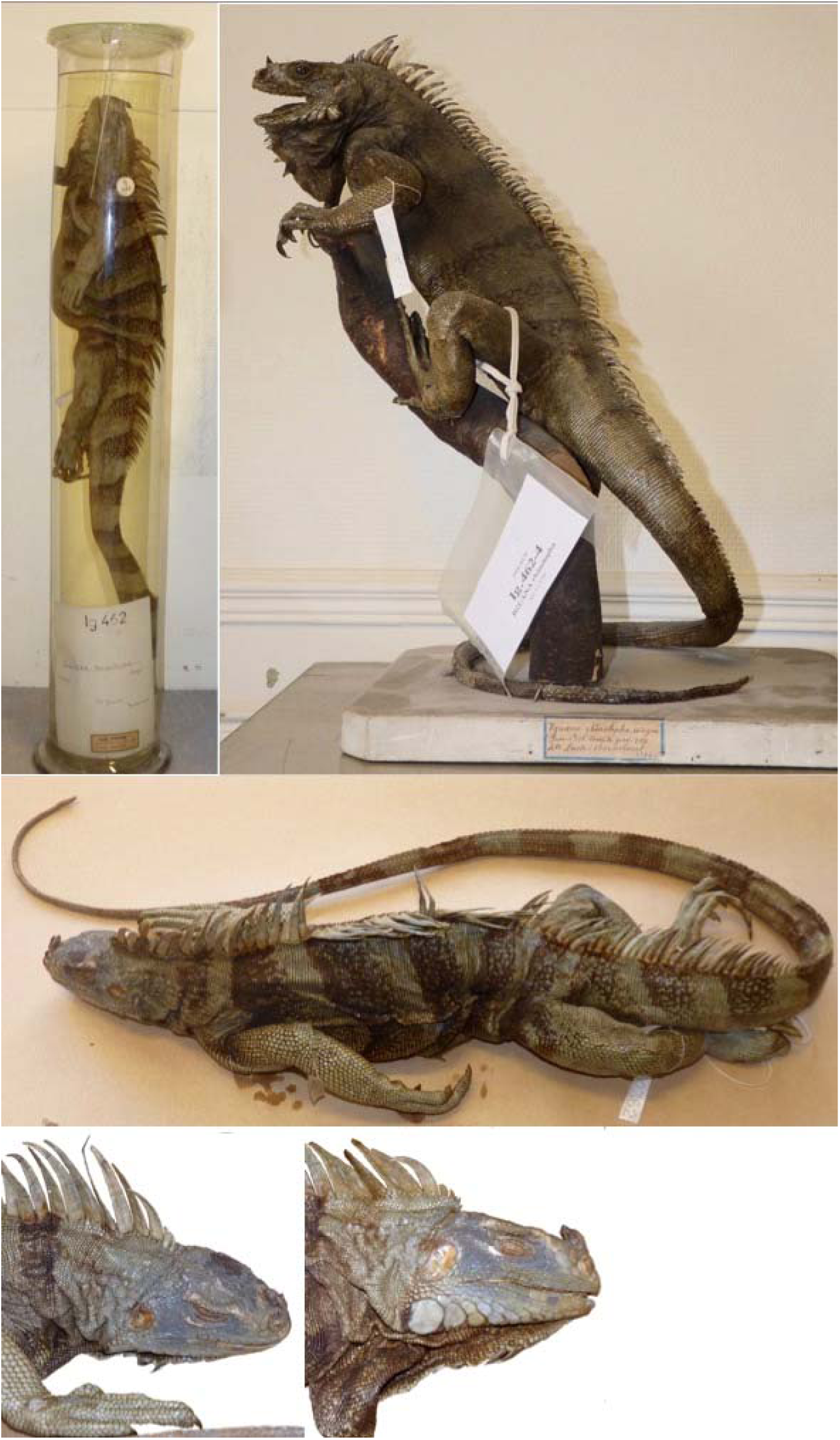
Holotype and paratype of *Iguana insularis sanctaluciae*. The holotype MNHN2362 is top left and the paratype MNHN 1996.8276 is top right. Both specimens collected by Bonnecour(t) in 1850-51 in Saint Lucia. Note the nasals (median and lateral horns), the small subtympanic plate, the low number of small nuchal tubercles, the 7 gular spikes, the prominent oval nostril, the banded body.

**FIGURE 8.**
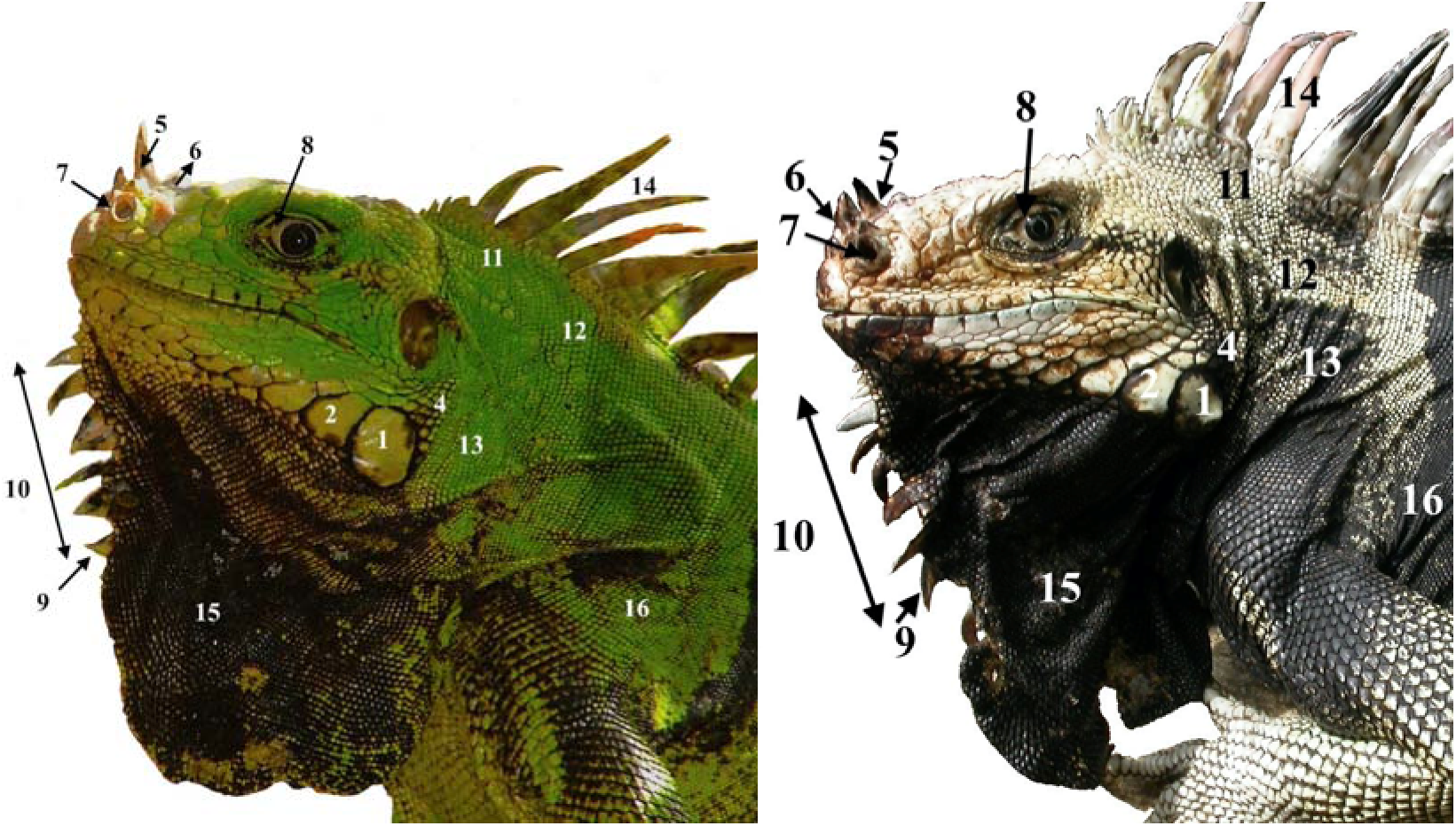
Ontogenetic change in colour of Iguana insularis sanctaluciae. Young male (left), old male (right). Annotations: 1. Small subtympanic plate ± 10% of the eardrum. 2. Two or three scales of decreasing size anterior to subtympanic plate. 3. Low number of sublabial scales with black margins. 4. No swelling of the jowls in breeding males. 5. Lateral and median horns. 6. Median horns with enlarged bases. 7. Oval to rounded nostril. 8. Brown eye with the white of the eye visible. 9. Triangular gular spikes. 10. 7 gular spikes. 11. Dispersed nuchal tubercles. 12. Low number of nuchal tubercles. 13. Small size of nuchal tubercles. 14. Orange in first dorsal spikes in breeding animals. 15. Entirely black dewlap in old adults. 16. Body and tail black and bright green in young individuals and very light green to almost pale greenish grey in old adults. Old individuals may look nearly “black and white”.

**FIGURE 9.**
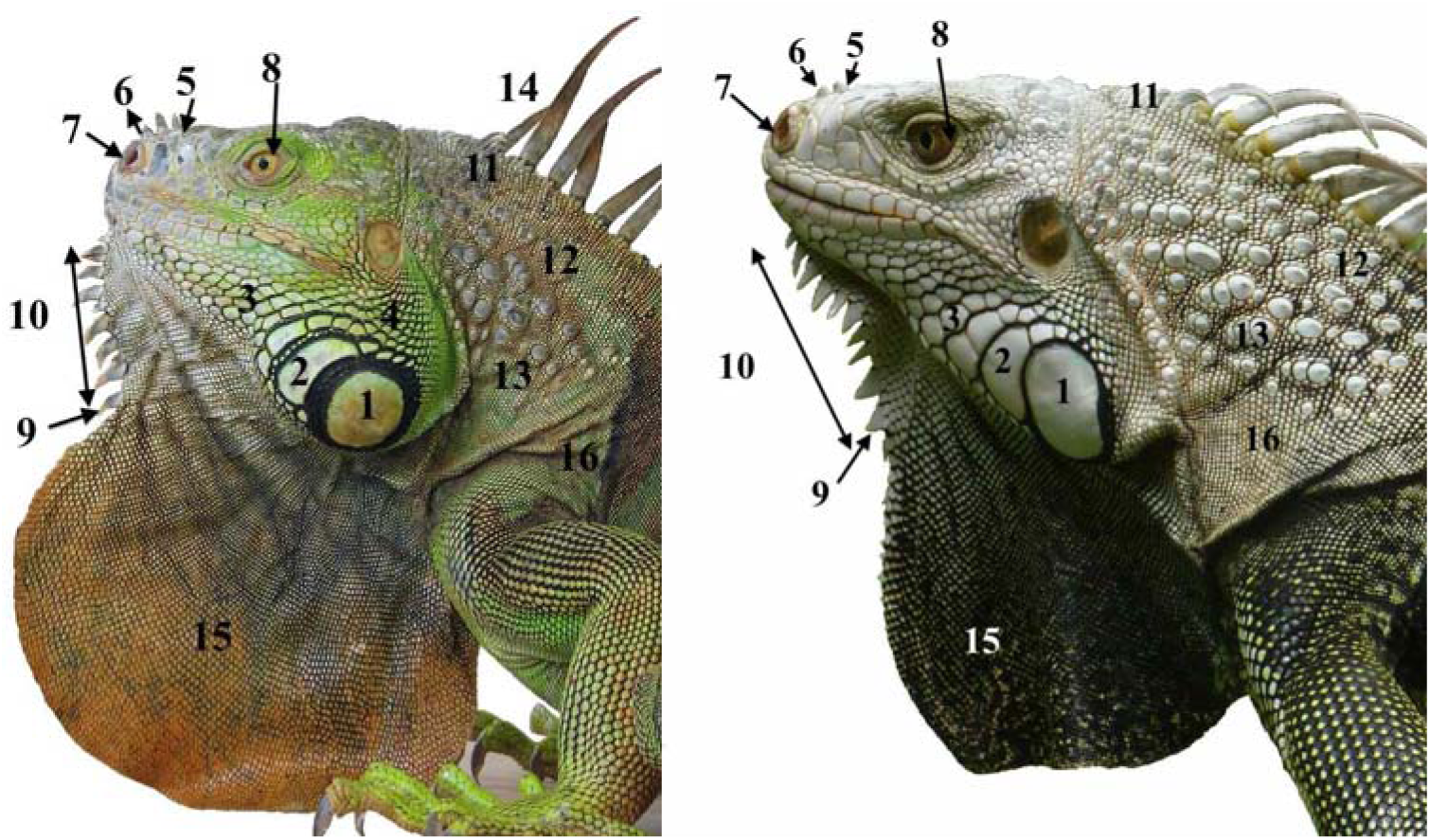
Horned iguanas from the Central American clade of the *“rhinolopha* phenotype. Photographed from invasive introduce populations in the Lesser Antilles: young male caught on Saint Maarten (left); old male caught in Saint Lucia (right). Annotations: 1. Huge subtympanic plate, 2 to 3 times the size of the eardrum. 2. A half crown of sublabial scales around the subtympanic plate and the first scale anterior to subtympanic plate. 3. Mosaic of sublabial scales. 4. Swelling of the jowls in breeding male. 5. Generally 2-3 small median horns and no lateral horns. 6. Flat small horns. 7. Triangular nostril. 8. Yellow to dark orange eye with not the white visible. 9. Triangular gular spikes. 10. Number of gular spikes ≥ 10. 11. Nuchal tubercles appear to be organised in rows. 12. High number of nuchal tubercles. 13. Very large nuchal tubercles. 14. Yellow, orange to red dorsal scales on the whole body in breeding males. 15. Variable size and colour of the dewlap but often large and not uniform black (cf. *I. insularis sanctaluciae*) or creamy white (cf. *I. insularis insularis*). 16. Body orange to red in breeding males, green in other individuals, and not heavily banded. This phenotype is recognised in this paper as a full species, *I. rhinolopha*, native to Central America (see text).

**FIGURE 10.**
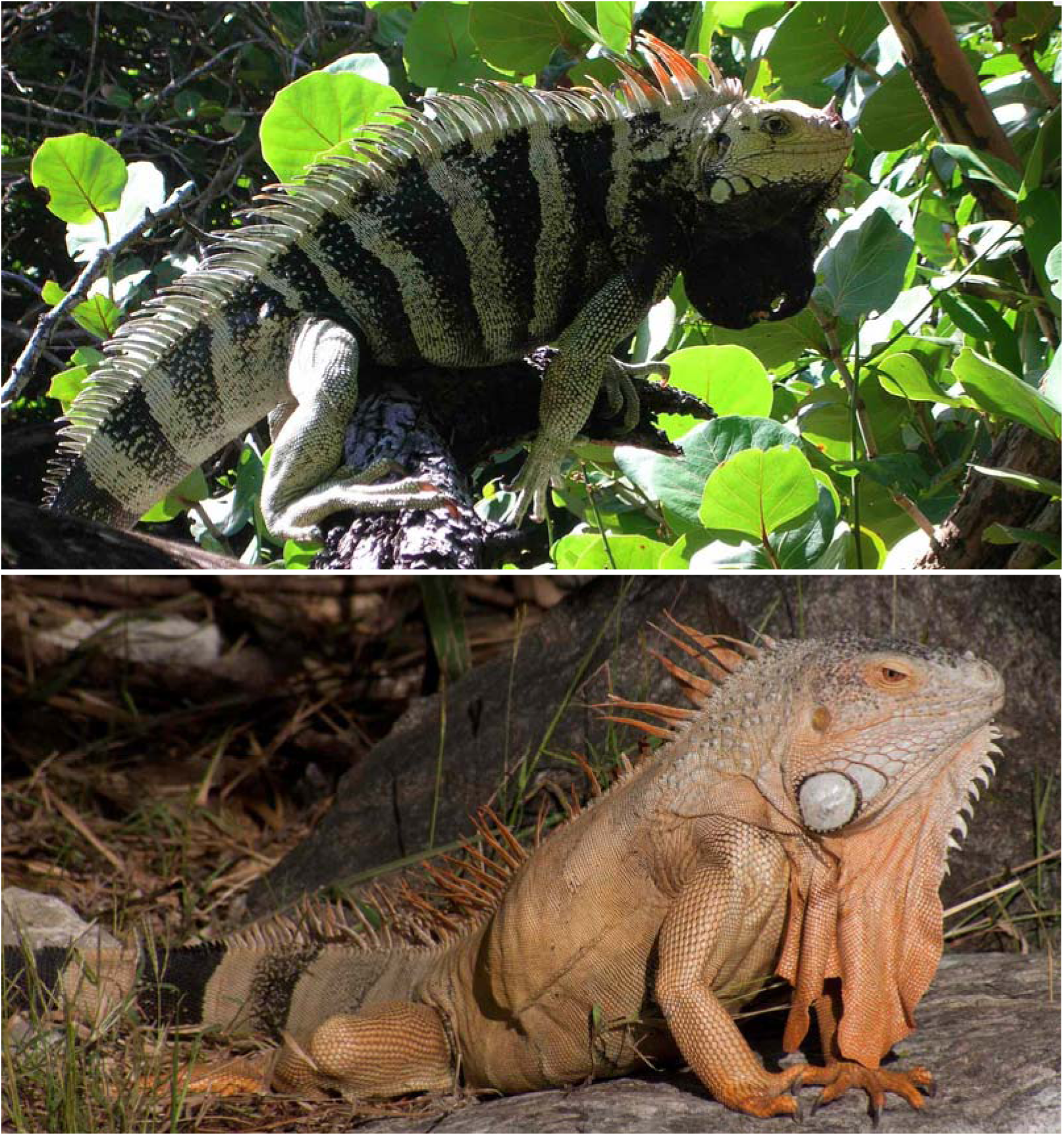
Adult breeding males *I. insularis sanctaluciae* and *I. rhinolopha*. The endemic Saint Lucia horned iguana (*I. insularis sanctaluciae*, photo from Grand Anse, top) is clearly different from the Central America horned iguana (*I. rhinolopha*: this specimen was photographed from an introduced population on Sint Maarten by M. Yokoyama, bottom) by size, body proportion, body colour, size and form of the horns, eye colouration, scalation of the jowls, and dewlap size, colour and form.

**FIGURE 11.**
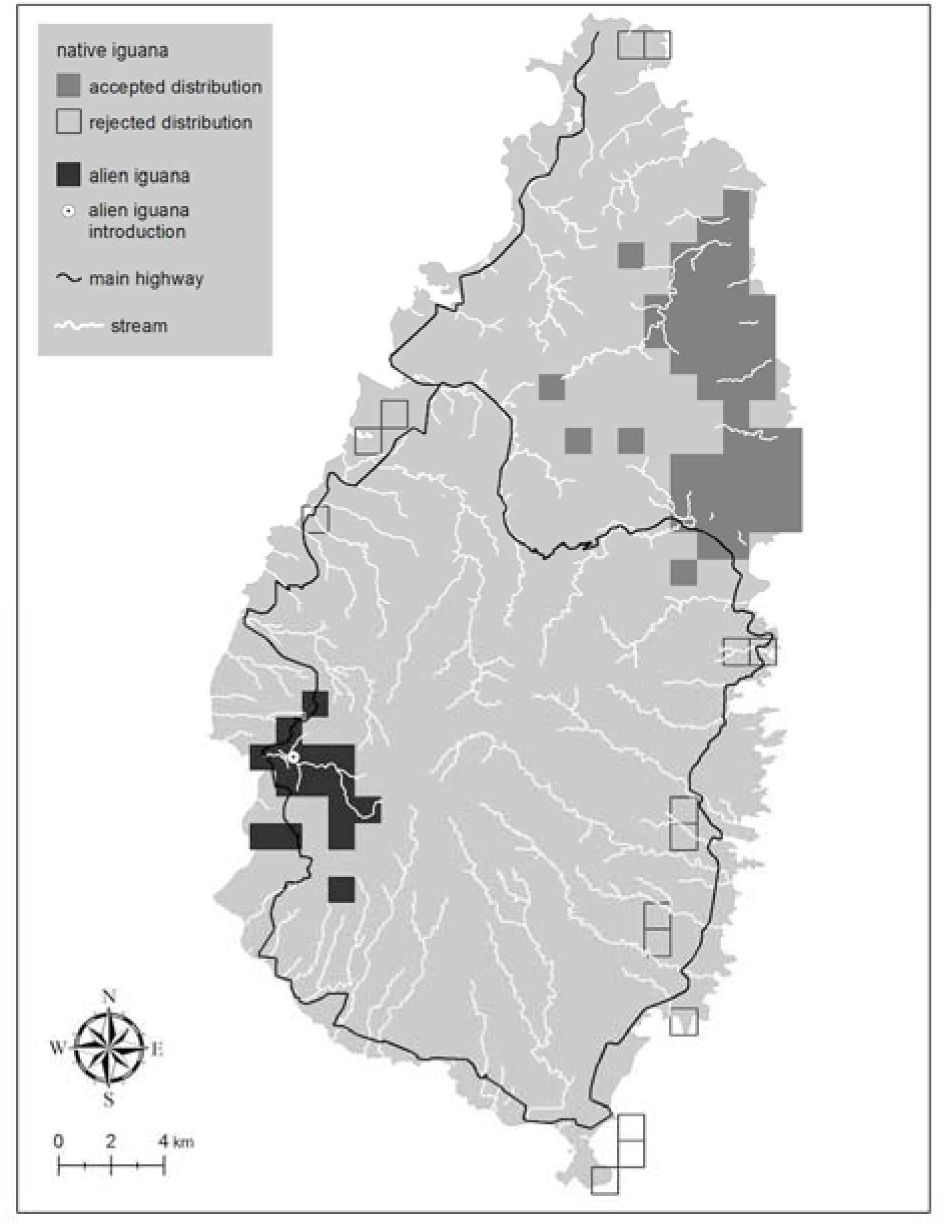
Distribution of iguanas on Saint Lucia. Locations are mapped to the nearest 1×1 km square. Data come from Morton *et al.* (2007; Saint Lucia). Black squares, Saint Lucia endemic iguana, Grey squares, alien iguana from Central America clade. “False absences” were minimized by interviewing persons about iguana presence in grid squares confirmed independently, through sightings and captures by us, to have iguanas present. We rejected some reported sightings of native iguanas, shown here as open grid squares, as being iguanas captured for food or pets or reports based on misidentification (for example on the islet of Maria Major off the far south of Saint Lucia; J. Lazell, *in litt.* 2010). Some of the reports that were accepted from the interior of the northern half of Saint Lucia may also be suspect, though they are all below 300 m ASL. These patterns of distribution suggest that the mountainous interior of Saint Lucia may create at least a partial barrier to direct east-west movements of iguanas.

#### Holotype

The holotype of *Iguana insularis sanctaluciae* housed in MNHN Paris under the number MNHN2362 and collected by Bonnecour(t) between 1850-1851. (Fig. 7)

This specimen is rigid, curved in its jar and it is nearly impossible to take accurate measurements.

***Sex*:** Male

***Age*:** Adult

***Morphological measurements*:** total length: 132 cm; SVL: 38.5 cm; tail length 93.5 cm; height and width of left subtympanic plate: 16.7/14.6; height and width of right subtympanic plate: 16.4/15.5, height of 4^th^ dorsal spike: 47.5 mm.

***Meristic characteristics*:** Number of gular spikes 7; Number of horns 2 median with very enlarged base + 3 small lateral on each side; Number of dorsal spikes to cloacae: 54

***Colouration*:** type in alcohol with discolouration, the ground colouration is green light grey with dark banding, 6 on the body and 10 on the tail, the scale of the dewlap are dark or half dark, the dorsal spikes are ochre but seem to have lost their original colour.

***Type locality*:** Saint Lucia, West Indies. No more information is known for this individual.

#### Paratype

The stuffed specimen MNHN 1996.8276 (Fig. 7) from the same island (Saint Lucia) and the same collector.

#### Diagnosis of *Iguana insularis sanctaluciae* (Figs 7-9)

*Iguana insularis sanctaluciae* resembles *I. insularis insularis*, but differs by the following association of characters:

– the scales of the jowls sometimes overlap;
– there are 7 or fewer triangular gular spikes of moderate size (cf. 8 or 9 exceptionally 10 gular spikes in *I. insularis insularis*);
– the vertical bands on the body are thicker, black and remain well developed in old individuals (cf. narrow bands on the body that fade with age in *I. insularis insularis*);
– the dewlap is black in old individuals (cf. creamy white in *I. insularis insularis*);
– the subtympanic plate and the associated 2-3 anterior scales have black pigmentation on their margins;
– Only the anterior dorsal spikes are orange in males (cf. most dorsal spikes have an orange hue in *I. insularis insularis*).

#### Size

The largest adult male to be measured on Saint Lucia was 160 cm in length (50 cm SVL) and weighed over 5 kg (Fig. 10). A sample of 30 adults in Saint Lucia had a mean total length of 110 cm (30 cm SVL) and mass of 1.3 kg.

#### Geographical distribution (Fig. 11)

The distribution of native Saint Lucia horned iguanas (*Iguana insularis sanctaluciae* subsp. nov.) and introduced alien iguanas (*Iguana rhinolopha*) on Saint Lucia is shown in (Fig. 16), redrawn from Morton & Krauss (2011) with minor updates, after an island-wide, systematic survey (Morton *et al.* 2007).

**Etymology:** The subspecific name is given in reference to Saint Lucia which is the only island inhabited by this new taxon.

#### Comparison to other species

*Iguana insularis* sp. nov. is distinguished from *I. iguana*, *I. rhinolopha* (considered here as a full species, see below) and *I. delicatissima* by the following combination of characters. Because in the field there is greatest risk of confusing the new species with invasive alien *I. rhinolopha*, which also has nasal horns, figures 9 and 10 highlight the main morphological differences between the anterior parts of *I. insularis* and *I. rhinolopha*.

#### Colour hue and pattern

The head, body and tail are bright green in young individuals, becoming very pale greenish grey or creamy white with age (unlike *I. iguana*, *I. rhinolopha* and *I. delicatissima*, which vary widely in hue but are rarely as pale). The body has 6-8 thin or thick vertical black bands (except in old adult *I. insularis insularis*, in which only faint traces of the vertical bands remain). These vertical black bands are present on the newborn *I. i. sanctaluciae* whereas they are generally absent in newborn *Iguana iguana* and *iguana rhinolopha.* According to Henderson & Powell (2018), juveniles most frequently are uniform green but this point has to be checked for *Iguana insularis insularis.* The tail has black bands that are conspicuous at all ages (unlike *I. delicatissima*, which does not have vertical bands on the body or tail). The legs are not black even in old individuals (unlike the indigenous iguanas of Saba and Montserrat). Although the body of pale adults may have a pinkish hue, the body colouration of breeding males is never orange as in *I. rhinolopha* from Central America (Fig. 10).

With age the dewlap changes from green to entirely creamy white (*I. insularis insularis*) or completely black (*I. insularis sanctaluciae*) as the indigenous iguanas of Saba and Montserrat, but never orange (cf. *I. rhinolopha*, Fig. 9). The dorsal crest is often high, especially in males (unlike *I. delicatissima*), and of the same colour as the light part of the body and often pink-orange towards the tips. The iris is dark brown, not yellow to orange, and the white part of the eye is visible (unlike *I. iguana*, *I. rhinolopha* and *I. delicatissima*). There is no black patch between the eye and the tympanum, and no pink on the jowls but some breeding males have pale golden yellow on the jowls.

#### Scalation

Several scales between the nostrils are elongated to form horns (whereas nasal horns are absent from *I. delicatissima* and *I. iguana*). There are 2 to 5 horns (usually 3-4) on the axial plane, and 1 to 3 smaller horns and sometimes up to 6 for *I. iguana insularis* on each side in adults (whereas lateral horns absent in *I. rhinolopha).* The horns are broad at their bases and the tallest are sometimes curved back (whereas the horns of *I. rhinolopha* are thin, straight and shorter). However, hatchlings and young juveniles of the new species have only very small horns.

The nostrils are prominent; their openings are from oval to circular, sometimes triangular in *Iguana insularis insularis*, looking from the side. There are some small to rarely medium conical scales on the occiput. There are 6-10 medium-sized gular spikes on the dewlap that extend to the half lower part. In adults, these spikes are triangular. A subtympanic plate is present (cf. absent in *I. delicatissima*) but it is relatively small: even in old adults the diameter of the subtympanic plate is no more than ± 20% the height of the tympanum (cf. 2-3 times the size of the tympanum in *I. rhinolopha*).

There are 2-3 scales of decreasing size anterior to the subtympanic plate, a characteristic not found in other species systematically present in *I. i. sanctaluciae* and sometimes in *I. i. insularis.* (This trait however resembles a feature of F1 hybrids between *I. iguana* and *I. delicatissima:* Breuil 2013, 2016). There are only few tubercular nape scales: fewer than 10 in *I. i. sanctaluciae* and up to 20 in *I. i. insularis*, small, not very prominent and dispersed, *i.e.* not arranged in more or less conspicuous rows as in *I. rhinolopha* (Fig. 10) and the largely melanistic iguanas of Saba and Montserrat populations (*I.* cf. *iguana*). This distinguishing character is present in hatchlings and throughout life, unlike some of secondary sex characteristics noted above.

#### Head

The head is relatively short and flat, and the dewlap is of medium size (cf. the large dewlap in *I. rhinolopha*). The scales anterior to the subtympanic plate overlap slightly in some individuals. The jowls do not appear swollen even in reproductively active males (cf. very well-developed jowls in breeding male *I. rhinolopha*).

## Results of genetic analysis

### Phylogeny

The intra- and interspecific relationships among *Iguana* species are shown in the phylogram (Fig. 12) and in the Median-Joining haplotype network (Fig. 13). Four clades can be observed:

**FIGURE 12.**
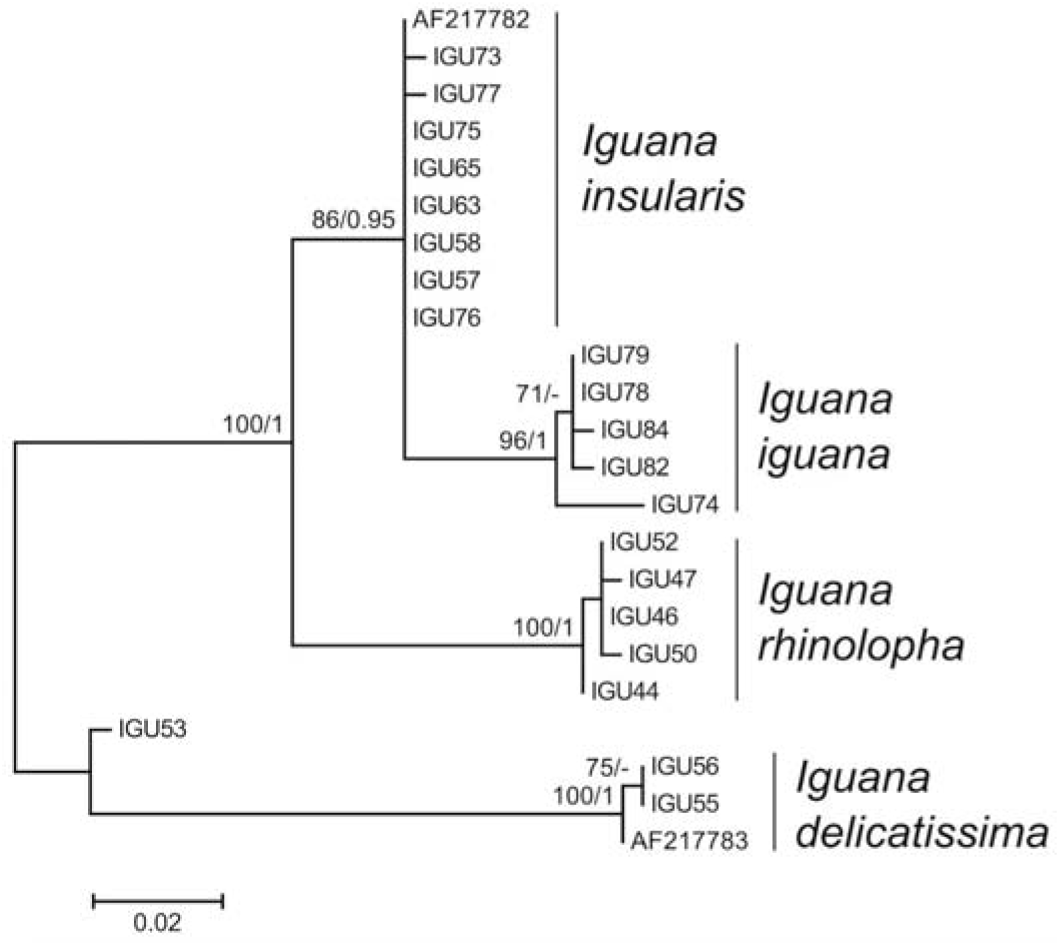
Phylogenetic tree. Based on mtDNA of 23 iguanas (21 from this study, 2 from GenBank). Four clades are identified. *Iguana delicatissima* (AF217783) serves as the outgroup. The monophyly of Lazell’s Southern Lesser Antilles group, characterised by horns, is described here as a new species *Iguana insularis.* The horned iguanas from Central America are also considered here as a full species *I. rhinolopha.* The sister group of *I. insularis* is *I. iguana* (based on specimens shown here from French Guiana). This phylogenetic tree shows that *I. iguana* is present as an invasive alien species in the Grenadines (IGU74) and that there is *I. delicatissima* mitochondrial DNA in some samples of *I. insularis sanctaluciae.* The ML tree with the highest log likelihood is shown. Node supports were indicated by bootstrap values from ML (>70) and posterior probability from BI (>0.95).

**FIGURE 13.** Median-Joining haplotype network. Based on 23 mtDNA sequences of *Iguana* (21 from this study, 2 from GenBank). Black circles are median vectors that represent extinct or unsampled haplotypes. Numbers of mutational steps are indicated by hatch marks.

Clade 1, the most basal, corresponds to *I. delicatissima* (GB: AF217783), which clustered with mtDNA from two iguanas from Grand Anse in Saint Lucia that had been identified in the field as pure endemic iguanas (see Discussion). IGU53, from the same locality, shows a haplotype that was found nowhere else.

Clade 2 corresponds to the mtDNA haplotypes shared by the alien iguanas on Saint Lucia, identified by their phenotypes as iguanas from Central America, likely originating from the pet trade (Fig. 10). These haplotypes are close to those of the Central American clade of Stephen *et al.* (2013) and Vuillaume (2012). This clade forms the sister group of the iguanas from French Guiana (South America) and from the southern Lesser Antilles.

Clade 3 groups the common green iguana (*I. iguana*) from French Guiana but also contains a specimen known only from a piece of shed skin (IGU74) found on Union Island (St Vincent & the Grenadines).

Clade 4 is consistent with our morphological analysis and shows the existence of an endemic species that inhabits Saint Lucia and the Grenadines (Grenada Bank).

### Genetic diversity

No linkage disequilibrium was detected after Bonferroni correction (adjusted P-value threshold = 0.0004). Only 5 of the 64 population-locus combinations deviated significantly from Hardy-Weinberg expectations (adjusted P-value threshold after Bonferroni correction = 0.0008). These deviations occurred for population of individuals endemic of Saint Lucia and so seem to be inherent to it. All microsatellite loci were polymorphic with an allelic richness (AR) ranging from 1 to 3.552 and a genetic diversity (He) ranging from 0 to 0.821 across populations (Table 2). Moreover, based on allelic frequencies, individuals introduced in Saint Lucia and those coming from French Guiana revealed the presence of several private alleles suggesting a specific genetic signature and so populations well genetically differentiated.

**Table 2.** Genetic diversity parameters for each locus Expected heterozygosity (He); number of private alleles (PA); allelic richness (AR); and inbreeding coefficient (Fis) have been computed for each population and each locus using FSTAT ver. 2.9.3.2 software (Goudet 2001). In italics and bold: the Fis with significant departures from Hardy-Weinberg expectations (i.e. significantly different from 0; P<0.0008 after Bonferroni adjustment).

|  |  | Saint Lucia<br>Introduced<br>*Iguana*<br>*rhinolopha* | Saint Lucia<br>endemic<br>*Iguana*<br>*insularis*<br>*sanctaluciae* | Grenadines<br>endemic<br>*Iguana*<br>*insularis*<br>*insularis* | French Guiana<br>*Iguana iguana* | All populations |
| --- | --- | --- | --- | --- | --- | --- |
|  | n | 8 | 17 | 4 | 7 | 36 |
| L2 | He | 0 | 0.265 | 0 | 0.262 | 0.132 |
|  | PA | 0 | 0 | 0 | 0 | - |
|  | AR | 1 | 1.647 | 1 | 1.692 | 1.91 |
|  | Fis | NA | 0.778 | NA | -0.091 | 0.343 |
| L3 | He | 0.125 | 0 | 0 | 0 | 0.031 |
|  | PA | 1 | 0 | 0 | 0 | - |
|  | AR | 1.375 | 1 | 1 | 1 | 1.768 |
|  | Fis | 0 | NA | NA | NA | 0 |
| L5 | He | 0.690 | 0.272 | 0 | 0.524 | 0.371 |
|  | PA | 2 | 0 | 0 | 0 | - |
|  | AR | 2.874 | 1.779 | 1 | 1.99 | 2.738 |
|  | Fis | -0.448 | **0.351** | NA | -0.091 | -0.063 |
| L6 | He | 0.429 | 0.217 | 0 | 0.524 | 0.2925 |
|  | PA | 0 | 0 | 0 | 1 | - |
|  | AR | 1.930 | 1.559 | 1 | 1.972 | 2.299 |
|  | Fis | -0.333 | 0.458 | NA | 0.727 | 0.284 |
| L8 | He | 0.667 | 0 | 0 | 0.262 | 0.232 |
|  | PA | 1 | 0 | 0 | 1 | - |
|  | AR | 2 | 1 | 1 | 1.692 | 1.453 |
|  | Fis | 0.5 | NA | NA | -0.091 | 0.204 |
| L9 | He | 0.652 | 0.217 | 0.75 | 0.607 | 0.556 |
|  | PA | 0 | 0 | 0 | 1 | - |
|  | AR | 2.604 | 1.559 | 2.929 | 2.692 | 2.972 |
|  | Fis | 0.233 | 0.458 | 0.333 | -0.412 | 0.153 |
| L13 | He | 0 | 0.217 | 0.5 | 0 | 0.179 |
|  | PA | 0 | 0 | 0 | 0 | - |
|  | AR | 1 | 1.559 | 1.964 | 1 | 1.975 |
|  | Fis | NA | 0.458 | 1 | NA | 0.729 |
| L14 | He | 0.125 | 0.116 | 0 | 0.143 | 0.096 |
|  | PA | 0 | 1 | 0 | 1 | - |
|  | AR | 1.375 | 1.353 | 1 | 1.429 | 1.959 |
|  | Fis | 0 | -0.016 | NA | 0 | -0.005 |
| L15 | He | 0.393 | 0.059 | 0 | 0.679 | 0.28275 |
|  | PA | 1 | 0 | 0 | 1 | - |
|  | **AR** | 1.885 | 1.176 | 1 | 2.774 | 1.893 |
|  | **Fis** | -0.273 | 0 | NA | 0.158 | -0.038 |
| **L16** | **He** | 0.81 | 0 | 0.25 | 0.143 | 0.301 |
|  | **PA** | 2 | 0 | 1 | 0 | - |
|  | **AR** | 3.375 | 1 | 1.75 | 1.429 | 2.06 |
|  | **Fis** | 0.824 | NA | 0 | 0 | 0.275 |
| **L17** | **He** | 0.548 | 0.224 | 0 | 0.488 | 0.315 |
|  | **PA** | 1 | 0 | 0 | 2 | - |
|  | **AR** | 1.99 | 1.652 | 1 | 2.275 | 3.055 |
|  | **Fis** | 0.478 | **0.475** | NA | 0.415 | 0.456 |
| **L18** | **He** | 0.339 | 0 | 0 | 0.533 | 0.218 |
|  | **PA** | 1 | 0 | 0 | 1 | - |
|  | **AR** | 1.786 | 1 | 1 | 1.998 | 1.667 |
|  | **Fis** | 0.632 | NA | NA | -0.25 | 0.191 |
| **L19** | **He** | 0.125 | 0.272 | 0 | 0.524 | 0.230 |
|  | **PA** | 0 | 0 | 0 | 0 | - |
|  | **AR** | 1.375 | 1.779 | 1 | 1.995 | 2.439 |
|  | **Fis** | 0 | **0.351** | NA | -0.364 | -0.004 |
| **L20** | **He** | 0.571 | 0.224 | 0 | 0.655 | 0.3625 |
|  | **PA** | 1 | 0 | 0 | 3 | - |
|  | **AR** | 2.348 | 1.652 | 1 | 2.827 | 3.213 |
|  | **Fis** | -0.094 | **0.475** | NA | -0.091 | 0.097 |
| **L23** | **He** | 0.4 | 0.272 | 0 | 0.821 | 0.373 |
|  | **PA** | 0 | 0 | 0 | 3 | - |
|  | **AR** | 1.909 | 1.779 | 1 | 3.552 | 3.071 |
|  | **Fis** | -0.25 | **0.351** | NA | 0.304 | 0.135 |
| **L24** | **He** | 0 | 0 | 0 | 0 | 0 |
|  | **PA** | 0 | 0 | 0 | 0 | - |
|  | **AR** | 1 | 1 | 1 | 1 | 1 |
|  | **Fis** | NA | NA | NA | NA | NA |
| **All loci** | **He** | 0.367 | 0.147 | 0.094 | 0.385 | 0.248 |
|  | **PA** | 10 | 1 | 1 | 14 | - |
|  | **AR** | 1 | 1 | 1 | 1 | 1 |
|  | **Fis** | 0.156 | 0.425 | 0.5 | 0.034 | 0.279 |

### Genetic structure

Results revealed significant genetic differentiation between populations. After applying the Bonferroni correction (adjusted P-value threshold = 0.0083), significant F_ST_ values were found between several pairwise populations (mean F_ST_ value = 0.495) (Table 3). This significant variation was corroborated by both FCA (Fig. 14) and the Bayesian individuals-based approach. Indeed, based on the individuals’genotypes, FCA clearly distinguishes four different populations and, furthermore, shows the low genetic diversity within the native Saint Lucia population (only 7 circles are shown because many individuals from Louvet and one from Grand Anse had the same genotypes). The STRUCTURE and STRUCTURE HARVESTER software revealed a highest delta K value of 3 (Figs 15, 16). According to these results, individuals from introduced iguana populations in Saint Lucia and those from French Guiana were mainly assigned to the first and second genetic cluster respectively (clades 3 and 2 from the phylogram, Fig. 12). Individuals native to Saint Lucia and the Grenadine islands were mainly assigned to the third genetic cluster (clade 4 of the phylogram). However, we can also distinguish three individuals (IGU53, IGU55, IGU56) that showed intermediate admixture coefficients, even though they were assumed to be of pure native origin. These intermediate admixture coefficients (Fig. 16, Table 4) and FCA results (Fig. 14) suggest hybridization has occurred in Grand Anse, where three out of the four individuals sampled are considered as hybrids between endemic and introduced individuals from different clades. Moreover, IGU55 and IGU56 have *I. delicatissima* haplotypes for ND4, and IGU53 has a unique haplotype closely related to *I. delicatissima* (Fig. 13).

**Table 3.** Pairwise Fst values for each population comparison (below diagonal) and their significance level (above diagonal). P-value threshold is adjusted with the Bonferroni correction, P= 0.0083.

|  | **Saint Lucia**<br>**Introduced**<br>***Iguana rhinolopha*** | **Saint Lucia**<br>**endemic**<br>***Iguana insularis sanctaluciae*** | **Grenadines**<br>**endemic**<br>***Iguana insularis insularis*** | **French**<br>**Guiana**<br>***Iguana iguana*** |
| --- | --- | --- | --- | --- |
| **Introduced Saint Lucia** | - | 0.0083 | 0.1083 | 0.0667 |
| **Saint Lucia endemic** | **0.6846** | - | 0.0583 | 0.0083 |
| **Grenadines endemic** | 0.6448 | 0.1051 | - | 0.0083 |
| **French Guiana** | 0.5048 | **0.5472** | **0.4832** | - |

**FIGURE 14.**
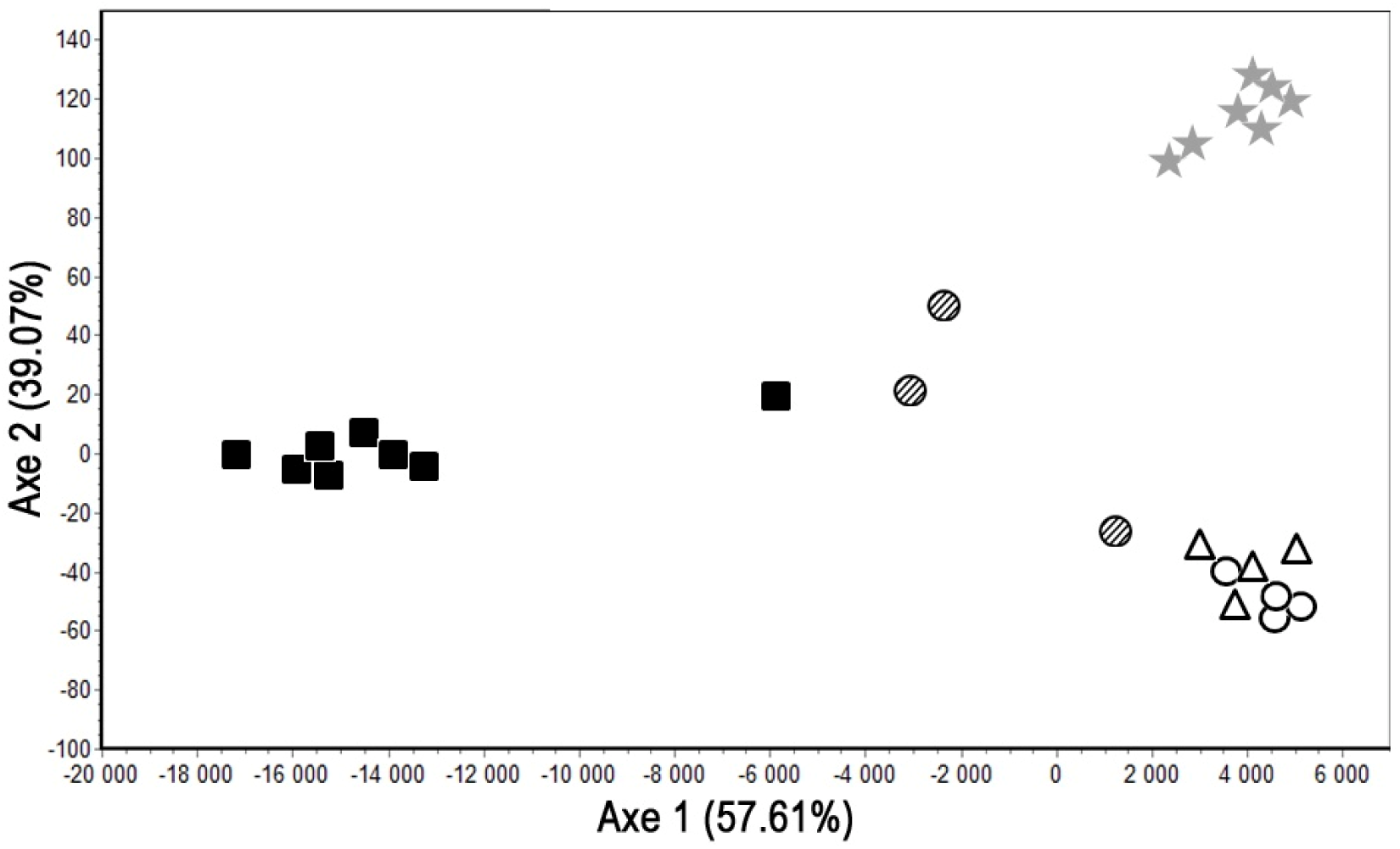
Factorial Correspondence Analysis (FCA) of the genotypes of samples from four populations. Black squares: individuals introduced in Saint Lucia (*Iguana rhinolopha*) [n=8] circles: individuals endemic to Saint Lucia (*Iguana insularis sanctaluciae*), with white circles for individuals from Louvet (n=13) and hatched circles for individuals from Grand Anse (n=4); white triangles: individuals endemic to Grenadines (*Iguana insularis insularis*), [n=4]; and grey stars: individuals from French Guiana (*Iguana iguana*) [n=7]. Only 7 circles are shown for the 17 ‘native’ individuals from Saint Lucia because many individuals from Louvet and one from Grand Anse had the same genotypes.

**FIGURE 15:**
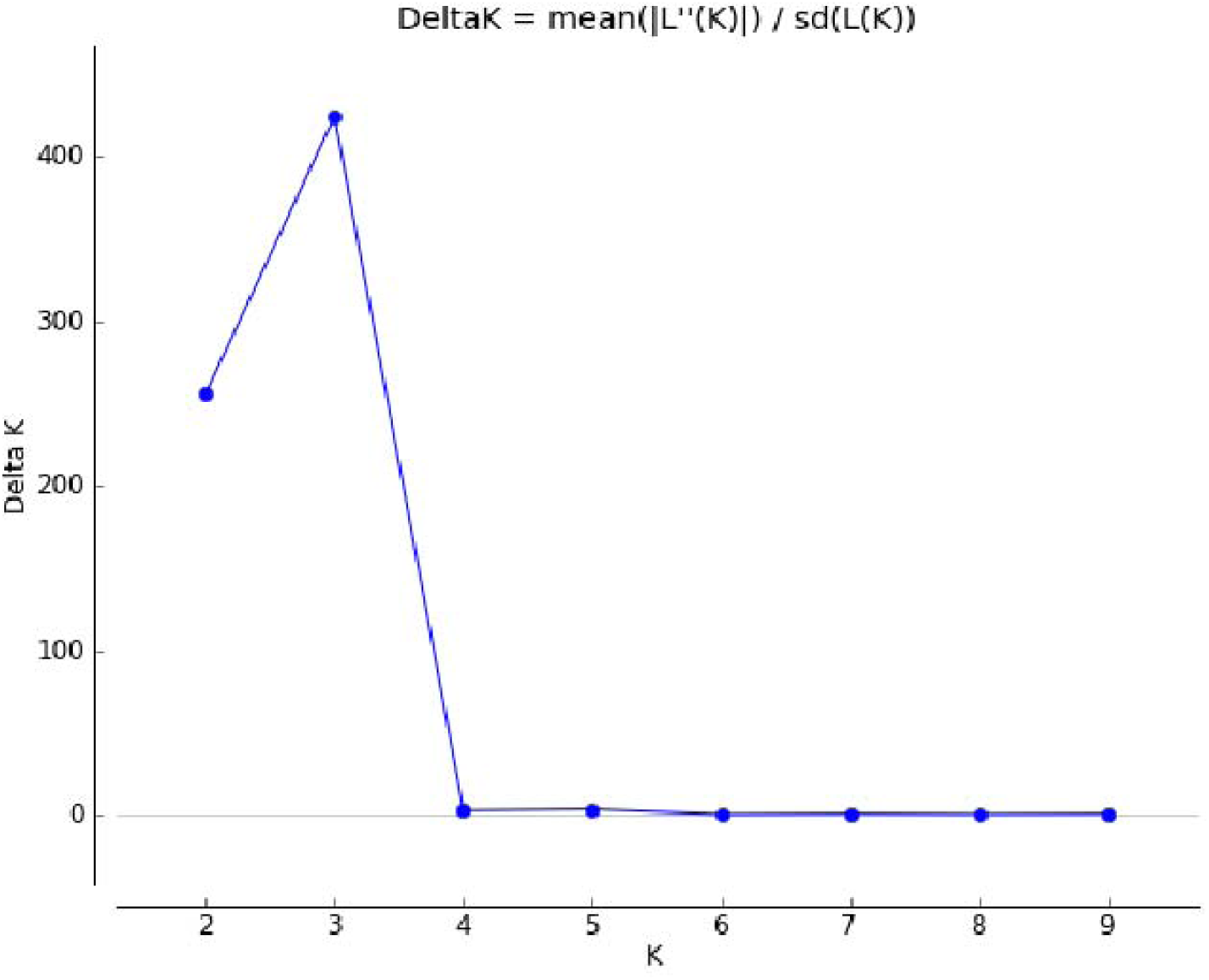
Results from the Evanno’s Method using the STRUCTURE HARVESTER software. This analysis reveals a maximum likelihood for K=3.

**FIGURE 16.**
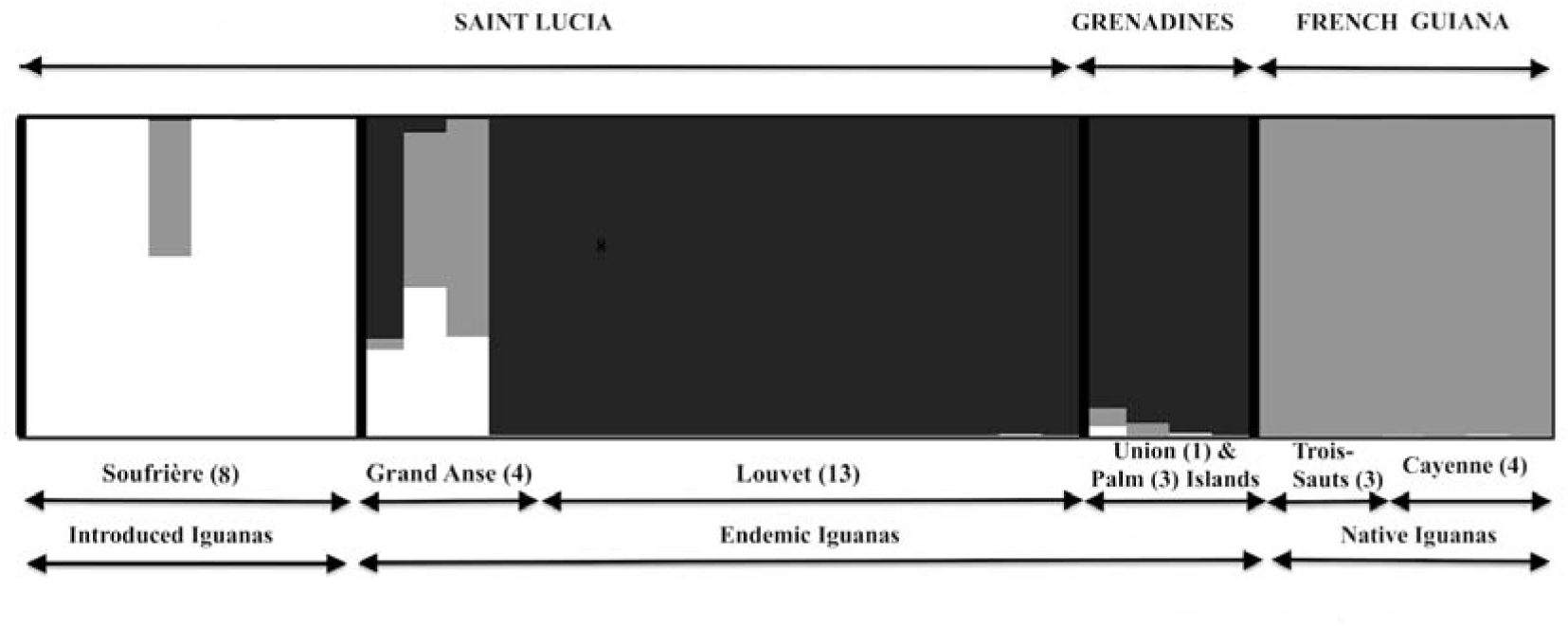
STRUCTURE bar plot. Showing admixture coefficient of each individual to the three inferred genetic clusters. This bar plot was produced using the DISTRUCT program (Rosenberg 2004).

**Table 4.** Admixture coefficient inferred by STRUCTURE software for the four studied populations.

|  | **Cluster 1** | **Cluster 2** | **Cluster 3** |
| --- | --- | --- | --- |
| **Introduced Saint Lucia**<br>***Iguana rhinolopha*** | 0.941 ($\pm 0.153$) | 0.056 ($\pm 0.151$) | 0.003 ($\pm 0.002$) |
| **Saint Lucia endemic**<br>***Iguana insularis sanctaluciae*** | 0.064 ($\pm 0.142$) | 0.072 ($\pm 0.196$) | 0.864 ($\pm 0.324$) |
| **Grenadines endemic**<br>***Iguana insularis insularis*** | 0.011 ($\pm 0.015$) | 0.025 ($\pm 0.025$) | 0.964 ($\pm 0.037$) |
| **French Guiana**<br>***Iguana iguana*** | 0.003 ($\pm 0.001$) | 0.994 ($\pm 0.001$) | 0.003 ($\pm 0.001$) |

## Discussion

### Taxonomic and systematic implications

The presence of horns on the iguanas of Central America first prompted the description of *Iguana rhinolopha*, by Wiegmann (1834), based on specimens caught in Mexico, and Duméril & Duméril (1851) and Boulenger (1885) subsequently applied the same name to iguanas from Saint Lucia because of their horns. No other morphological traits were used to distinguish *I. rhinolopha* apart from a small difference in number and size of spikes on the dorsal crest, identified by Duméril & Bibron (1837) based on a very small number of specimens (Figs 9, 10). Later, with the widespread use of the subspecies concept, Dunn (1934) proposed that *rhinolopha* was merely a subspecies of *I. iguana*, and Barbour (1935) also followed this position. This was the consensus until the work of Lazell (1973), who realised that the presence of horns on the snout was inconsistent and, because it occurs polytopically, he rejected the taxon *rhinolopha.* Lazell realised that the indigenous iguanas on Saint Lucia possessed horns that were generally well developed on mature adult individuals, and this is also confirmed by our observations. However, while horns can be found in Central American iguanas, they differ from the arrangement of horns in *Iguana insularis* sp. nov., as we have demonstrated in this work.

This paper describes a new species of horned iguana, *Iguana insularis* sp. nov., known only from the southern Lesser Antilles. All the indigenous iguanas from Saint Lucia, St Vincent & the Grenadines and Grenada, described herein as *Iguana insularis*, possess median and lateral horns. Moreover *I. insularis* has a combination of morphological traits that makes it unique, including: low number and small size of nuchal tubercles, small subtympanic plate, brown iris colour with the white of the eye visible, no subtympanic swelling, colour hue and pattern of the body, etc. (Figs 4, 5, 7, 8). With about 2% divergence in the ND4-Leu sequence, *I. insularis* shows a level of differentiation from *I. iguana* from French Guiana that is within the interval of divergence of subspecies recognition among *Cyclura* (Malone & Davis 2004). So, should *insularis* be regarded as a full species, or simply a horned subspecies of *Iguana iguana*? In recent years, the subspecies concept has become unpopular in herpetology, and almost every subspecies in the Caribbean has now been upgraded to a full species (*e.g.* Breuil 2002) in accordance with the phylogenetic species concept that treats populations as separate species if they are on separate evolutionary trajectories (see *e.g.* Torstrom *et al.* 2014), as is usually the case for animals confined to separate islands. Aside from the presence of horns, we consider there are enough distinct and consistent morphological and genetical differences to recognise the native southern Lesser Antillean iguanas as a full species, *I. insularis*.

While the native iguanas of the Southern Lesser Antilles undoubtedly share a common ancestry (Figs 12, 13, 14, 15), consistently strong differences in colour hue and pattern have prompted us to describe the endemic iguanas of Saint Lucia as *I. insularis sanctaluciae* and those of the Grenada Bank as *I. insularis insularis.* The same basis was used by Hawlittsckek *et al.* (2012) for distinguishing species and subspecies of Comoran snakes. Given that the two new subspecies are reproductively isolated on separate island banks, we cannot discount the possibility that further investigation may lead to them being further elevated into separate species. Further research is also needed on the island of St Vincent to determine whether any purebred (i.e. non-hybrid) native iguanas remain here, and whether they belong to either of the aforementioned subspecies or a third, undescribed subspecies.

When did *Iguana insularis* diverge from other species in this genus? Studies of iguana morphology (Breuil 2013, 2016) and genetics (Stephen *et al.* 2013; Valette *et al.* 2012; Vuillaume *et al.* 2015) have shown there are at least three ancient lineages (*I. delicatissima* and the iguanas of Central America and South American) in the genus *Iguana*, with a genetic divergence approximated by a molecular clock of 1.29 million years for every 1% sequence divergence at the ND4-Leu Locus (Malone *et al.* 2000). With approximately 10% divergence (Malone & Davis, 2004) between *I. iguana* and *I. delicatissima*, the age of separation of these two species is therefore about 11-12 My; but according to Hedges *et al.* (2015), the two lineages could have diverged as much as 22.8 My ago. If we take the lower value of the molecular clock, *I. insularis* diverged about 2.2 My ago. This timeframe is compatible with the ages of Saint Lucia and Grenada, where the oldest rocks date from the Miocene and belong to the intermediate volcanic arc (Bouysse & Garrabé 1984; Germa 2008).

Evolution in isolation over millions of years does not automatically mean the taxa cannot interbreed. Even the most distantly related species in the genus *Iguana* – *I. delicatissima* and *I. iguana* – can interbreed to produce healthy, fertile offspring (Breuil 2013, 2016; Vuillaume *et al.* 2015). This is not very unusual among even more distantly related taxa: For example, even crocodiles in the genus *Crocodylus* from opposite sides of the globe can interbreed to produce fertile hybrids that have a competitive advantage (e.g. Daltry *et al.* 2016a). Judging from their nesting periods (Fig. 17), there is some overlap between the breeding seasons of *I. insularis*, *I. delicatissima* and introduced *I. iguana* from South America, which might enable these species to interbreed.

Where does this leave the horned iguanas of Central America? Stephen *et al.* (2013) recognized two well supported genetic groups in *Iguana iguana* as evolutionary significant units: Central America (México to Panamá) and South America (including Curaçao and the Lesser Antilles) but declined to propose any taxonomic changes pending further sampling across Panamá and South America and a better understanding of the basal position of the populations of Curacao (Buckley *et al.* 2016). The divergence between Central and South American clades based on ND4-Leu is about 4.3%, similar to the 4% divergence between *Cyclura* species (Malone & Davis 2004). We therefore accept *I. rhinolopha* (Wiegmann 1834) as a full species native to Central America, distinguished not only by its unique arrangement of nasal horns but numerous other morphological and genetical characters, as shown by Breuil (2013, 2016), Stephen *et al.* (2013) and Vuillaume *et al.* (2015). This distinctive clade forms a basal lineage with respect to *I. iguana* from South America and the iguanas of the Lesser Antilles (Fig. 13). By sampling more areas across this region, it is likely other new cryptic species will be identified (Bickford *et al.*, 2007; Buckley *et al.* 2016). Indeed, the melanistic iguanas from Saba and Montserrat have been shown to be morphologically and genetically distinct (Breuil 2013, 2016; Stephen *et al.* 2013; Vuillaume *et al.* 2015) and will be considered in another work. Nevertheless, further studies elsewhere in this region are unlikely to change our conclusions regarding the relationships of *Iguana insularis* vs *I. rhinolopha* and *I. iguana* in the Lesser Antilles.

The most unexpected revelation from our genetic study is that two iguanas (IGU55, IGU56) from Grand Anse, Northeast Saint Lucia, had *delicatissima* ND4 haplotypes, while a third (IGU53) had an unknown haplotype that also clustered with *delicatissima* in addition to the microsatellites from both *I. iguana* and *I. rhinolopha* (Figs 12, 13, 16). The samples used to characterise the population of Grand Anse were juveniles caught 10 years ago, when the morphological differences between *Iguana insularis*, *I. iguana*, *I. rhinolopha* and *I. delicatissima* were less well understood, and the animals were erroneously registered as pure native Saint Lucia iguanas (Table 1). As reported in our introduction, Provencher (1880) had mentioned the presence of *I. delicatissima* on Saint Lucia, but the only evidence was a poorly executed drawing (reproduced in Fig. 1). Our discovery of *Iguana delicatissima* haplotypes in Saint Lucia may be explained by either an ancient *delicatissima* population on Saint Lucia that persists only as a maternal lineage or by the recent arrival of a female with an unknown haplotype from another island that has reproduced with *Iguana insularis sanctaluciae.* Currently, however, we have insufficient data to support one or other hypothesis.

### Conservation status of *Iguana insularis* sp. nov

The new species of iguana is at risk throughout its range in the southern Lesser Antilles due to habitat loss, hunting (both for bushmeat and the pet trade) and invasive alien species, especially alien predators and non-native *Iguana* species.

Of the two new subspecies described in this paper, the Saint Lucia horned iguana (*I. insularis sanctaluciae*) appears to be most scarce and vulnerable to extinction. These iguanas are a fully protected species under the Wildlife Protection Act (Laws of Saint Lucia 2010), but they continue to be hunted and eaten at a significant level (Morton & Haynes, pers. obs.) and have been illegally exported and sold to collectors overseas (J. Daltry, pers. obs.). The range of the native iguana population on Saint Lucia is now restricted to that part of the island without good road access (Fig. 16), putatively because of over-hunting in the more accessible areas.

Habitat conversion for development (in particular the proposed tourism developments on the three large estates of Louvet, Grand Anse and Marquis, and the proposed North East Corridor highway) is currently considered the most severe threat facing the remaining population. Illegal mining of beach sand threatens the iguanas’ nesting sites and their seasonal deciduous forest habitat is also especially vulnerable to wildfires (Robbins *et al.* 2008). Developments in the Northeast are also likely to exacerbate threats from introduced mammalian predators: Feral cats (*Felis catus*), southern opossums (*Didelphis marsupialis*) and small Asian mongooses (*Herpestes auropunctatus*) are all known to kill hatchling iguanas in Saint Lucia (Morton *et al.* 2007). Mongooses also take iguana eggs, and the mutilation of hatchlings whilst still in the nest chamber has been attributed to rats (*Rattus rattus* or *R. norvegicus*) (Morton pers. obs.). Both feral and domestic dogs (*Canis familiaris*) prey on iguanas, especially nesting females that are especially vulnerable whilst on the ground (Morton *et al.* 2007).

Invasive alien iguanas also pose a serious threat, having become well established in Southwest Saint Lucia. Despite numerous efforts to catch and cull the invasive iguanas (Morton & Krauss 2011; Krauss 2013; Krauss *et al.* 2014), it is proving prohibitively expensive and difficult to limit their spread and prevent contact with the indigenous *Iguana insularis* in Northeast Saint Lucia (Fig. 16), especially with the risk of human-mediated transport across the country. The alien iguanas (*Iguana rhinolopha*) appear to originate from the Central America clade, characterised by its greatly enlarged subtympanic plate, yellow iris, numerous and conspicuous tubercular nape scales, orange colouration in breeding male, huge dewlap with more than ten spikes but also small horns on the stout (Breuil 2013, 2016). The invasive iguanas on Saint Lucia (Fig. 9) also have larger clutches than the native species: mean clutch size for the former is 40 (n = 4 clutches) and mean clutch size for *I. insularis sanctaluciae* is only 23 eggs (n = 14 clutches). The same invasive iguanas from Central America have clutch sizes of 8-75 in Puerto Rico (Lopez-Torres *et al.* 2012) and 20–63 in Hawaii (McKeon 1996). It is very likely that the alien iguanas could hybridize as suggested by the genetic analysis (Fig. 17) with and outcompete *Iguana insularis sanctaluciae*, leading to its elimination (as occurred with *I. delicatissima* in Les Saintes, Basse-Terre and Grande-Terre: Breuil 2002, 2013, 2016; Vuillaume *et al.* 2015).

**FIGURE 17.**
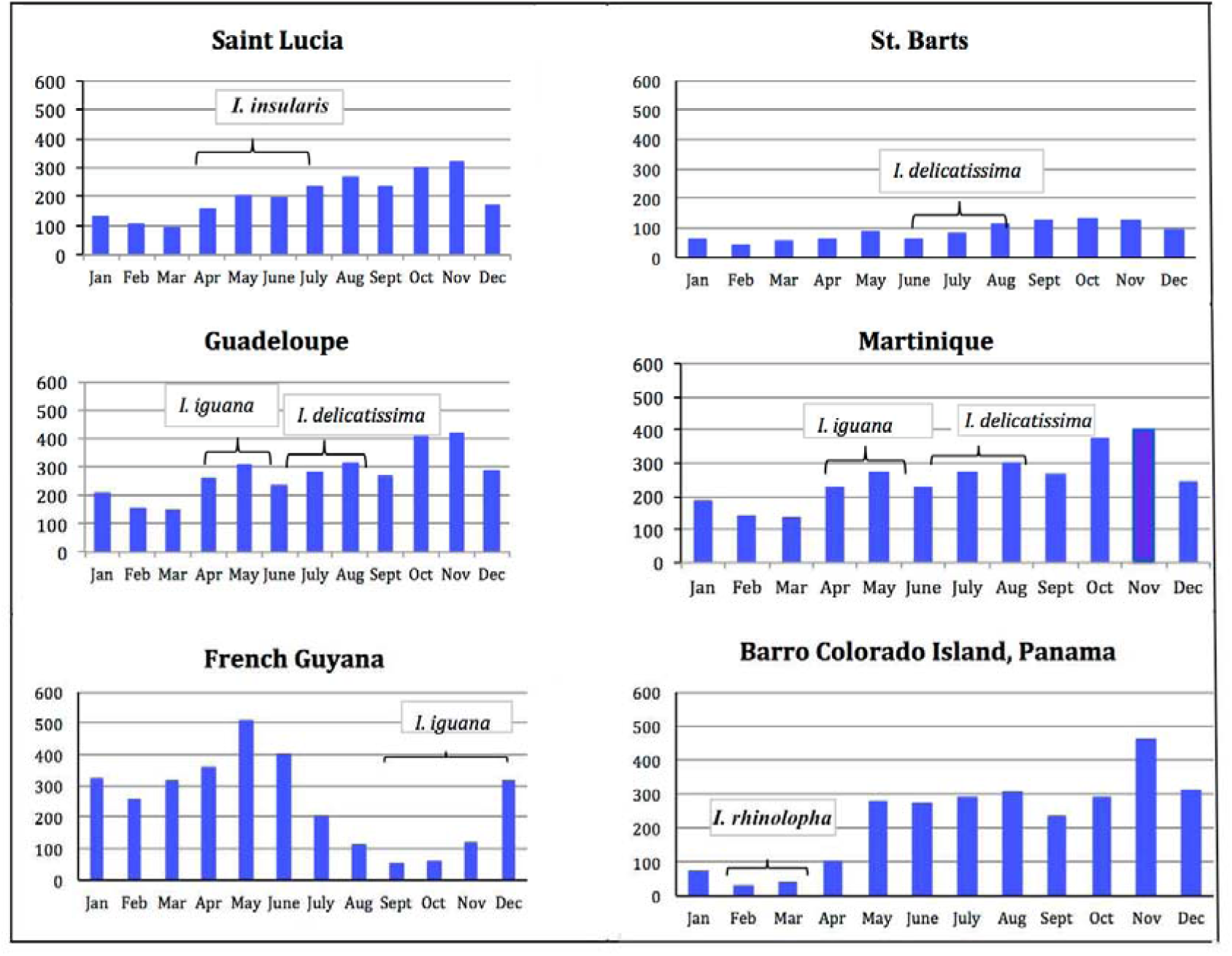
Nesting periods of various iguana populations. Monthly mean precipitation is indicated in mm. Precipitation varies according to altitude and aspect, but the important point is that after a 3 months incubation period, the eclosions occur at the beginning or during the rainy season. The brackets indicate the main laying period for the different species at different insular and continental locations. Note that the nesting period of *Iguana insularis sanctaluciae* overlaps the laying periods of both *Iguana delicatissima* and *Iguana iguana*, which suggests that mating periods could also overlap, potentially enabling all three species to interbreed.

The status and threats to the Grenada Bank subspecies is less well understood because iguanas have been less closely studied and existing literature on iguanas has failed to distinguish between the native and invasive alien iguanas. Given that iguanas in general are considered to be fairly abundant and widespread on Grenada and St Vincent & the Grenadines, adult iguanas may still be lawfully hunted for several months of the year (typically October through December or January) for personal consumption and local sale (Laws of Saint Vincent & the Grenadines 1990; Laws of Grenada 1990). Unlike Saint Lucia, the national laws here do not distinguish between native and introduced or hybrid iguanas, nor define any populations that may not be hunted or moved within national borders. It is therefore not uncommon for hunters to collect iguanas from the Grenadine islands for sale on St Vincent or Grenada during the hunting season (G. Gaymes, pers. obs.). Hunting is frequent on the uninhabited island of Balliveaux, for example, where hunters from Bequia and Saint Vincent “carry away dozens of iguanas” (Daudin & Da Silva 2011). In this context of numerous translocations, it is uncertain how many purebred populations of *Iguana insularis insularis* remain.

Not surprisingly, considering the lack of any concerted effort to prevent incursions, alien iguanas appear to have become very widespread in Grenada and St Vincent & the Grenadines, with perhaps no purebred (i.e. non-hybrid) native iguanas remaining anywhere on the main islands of St Vincent or Grenada. We suspect that the most intact native populations are restricted to some of the smallest islands in the Grenadines, including Palm Island and Union Island, where genetic samples were analysed for this study. It is noteworthy that in this context that IGU74 collected as a shed skin has a ND4 haplotype that clusters with *Iguana iguana* from South America. Like the iguanas of Saint Lucia, even these populations are at substantial risk from invasive alien predators (dogs, cats, opossums, etc.) and coastal deforestation and development (Daltry *et al.* 2016b).

Under the national laws of all three countries – Saint Lucia, St Vincent & the Grenadines and Grenada – the export of iguanas or their products is prohibited without permits from their respective chief wildlife wardens. Furthermore, on the basis that the iguanas were classified as *Iguana iguana*, listed on CITES Appendix II, exports have required an export certificate from the CITES Management Authority. Nevertheless, recent years have seen a rise in young iguanas being smuggled from these islands and sold under various trade names, including the Saint Lucia iguana, pink rhino iguana (originating from Union Island) and white zebra rhino iguana (from the Tobago Cays). Iguanas from Saint Lucia and the Grenadines have been offered for sale in the USA, Japan and Europe, for prices of up to $10,000 per pair, with traders often claiming to have CITES permits, even though no such export permits have been issued by these countries (Noseworthy 2017).

At species level, *Iguana insularis* sp. nov. qualifies as Vulnerable, under IUCN Red List criteria B1a,b (i-v), B2 a,b (i-v) and probably C2a(i). Pure-bred (non-hybrid) individuals have been confirmed only in three locations (Northeast Saint Lucia and, in St Vincent & the Grenadines, on Union Island and Palm Island), giving a known extent of occurrence of less than 2,000 km^2^ and an area of occupancy of less than 30 km^2^. Further surveys and genetic analysis are needed to verify status, but it is unlikely there are as many as 10 locations with non-hybrid populations of *Iguana insularis*, and no population is known to contain more than 1,000 mature individuals. An ongoing decline is predicted due to introgressions from invasive alien iguanas, invasive alien predators, habitat loss, and over-collection for meat and the pet trade. (Note that the measured extent of occurrence includes marine areas between the islands: a common problem when applying this method to taxa on islands).

At subspecies level, the Saint Lucia iguana (*Iguana insularis sanctaluciae*) is Critically Endangered under criteria B1ab(i-iii): Its extent of occurrence is approximately 30 km^2^, it exists in only one location (Northeast Saint Lucia), and a continuing decline is observed and projected in the (i) extent of occurrence, (ii) area of occupancy, and (iii) area, extent and quality of habitat due to tourism developments, sand-mining, livestock grazing and other documented threats (Daltry 2009). The Grenadines horned iguana (*Iguana insularis insularis*) has been less closely studied but qualifies as Vulnerable and possibly Endangered under criteria B1ab(i-iii): Its area of occupancy is less than 20 km^2^, it exists at not more than 10 locations (only two locations – the 9 km^2^ Union Island and 0.55 km^2^ Palm Island – have been confirmed to have reasonably intact, non-hybrid populations) and estimates indicate continuing decline, observed, inferred or projected, in the (i) extent of occurrence, (ii) area of occupancy and (iii) area, extent and quality of habitat due to tourism developments, livestock grazing, bushfires and other threats (Daltry *et al.* 2016).

### Recommendations

International trade in horned iguanas from the Grenadine islands (specifically, Union Island, Palm Island and the Tobago Cays) and Saint Lucia has been confirmed in recent years at a level that could present a serious risk to both subspecies. By formally naming the new species and the two subspecies, we recognise that the demand from reptile collectors could increase (Auliya *et al.* 2016). We therefore recommend that as an urgent precaution the species *Iguana insularis* sp. nov., should be placed on Appendix I of CITES at the next Conference of Parties to monitor and control illegal international trade. As an interim measure, we urge Saint Lucia, Saint Vincent & the Grenadines and Grenada to jointly request the CITES Secretariat to place *Iguana insularis* on Appendix III. This is necessary to help to ensure that iguanas cannot be sold overseas without a CITES export permit from the country of origin.

Nationally, the new species is fully protected only in Saint Lucia, where the Wildlife Protection Act distinguishes between the native Saint Lucia iguana and the (non-protected) invasive alien iguanas. We recommend Grenada and St Vincent & the Grenadines also consider increased levels of protection for the native horned iguanas and ensure that any future exploitation of *I. insularis* populations is monitored closely and sustainable. Considering the outstanding importance of the apparently purebred and growing population of *I. insularis insularis* on Palm Island (St Vincent & the Grenadines), technical assistance should be offered to the landowners to find solutions to complaints that the iguanas are causing a nuisance.

Because invasive alien iguanas have already become well established in all three countries, it is also imperative to safeguard all remaining native iguana populations from hybridisation and competition. Active biosecurity measures must be developed to prevent non-native iguanas from successfully spreading to Northeast Saint Lucia, Palm Island, Union Island and any other areas known to have purebred (non-hybrid) *I. insularis*, including monitoring these native populations regularly to ensure any incursions are detected and dealt with swiftly. Further surveys are required on St Vincent and Grenada to determine whether any purebred native iguanas remain on these islands. If alien iguanas continue to increase unchecked on Saint Lucia, it may be necessary to separate them from the native iguanas with a physical barrier. With this in mind, plans are currently being developed to create a ‘mainland island’ sanctuary for native Saint Lucian wildlife, surrounded by a pest-proof fence (Saint Lucia Forests and Land Resources Department, 2015) that could potentially conserve several hundred *I. insularis sanctaluciae* in strict isolation from alien iguanas.

To support all these recommendations, it will be necessary to develop illustrated identification materials for researchers, enforcement officials and other stakeholders to reliably distinguish both subspecies of *I. insularis* at all ages from *I. iguana*, *I. rhinolopha* and other species.

Thus, we hope that the recognition of this new species and its two subspecies will ultimately facilitate their protection and conservation.

## Acknowledgements

This project was financed mainly by the Direction Régionale de l’Environnement et du Logement de Martinique to study the origin of invasive common green iguanas in Martinique and their relationships to the Saint Lucia iguana. The genetic study formed part of Barbara Vuillaume’s Master’s degree. The genetic analyses were initially conducted at Genindexe in La Rochelle (France) and completed by David Schikorski in Genindexe Labofarm (France), which financed part of this study.

Specimens were collected from Palm Island and Union Island, in the Grenadines, by JD and GG with kind permission from the Palm Island Resort and the St Vincent & the Grenadines Forestry Department. Fieldwork was supported with funding from the FFI Species Fund, Disney Conservation Fund, National Geographic, US Fish & Wildlife Service (#F18AP00796), and the St Vincent & the Grenadines Preservation Fund. We are most grateful to Roseman Adams, Katrina Adams and other members of the Union Island Environmental Attackers for their assistance. Special thanks also to F. Catzefis (CNRS, Montpellier) and Benoît de Thoisy (Institut Pasteur, Cayenne, French Guiana), who provided samples from French Guiana.

All Saint Lucia samples were collected by the Durrell Wildlife Conservation Trust and the Saint Lucia Forestry Department. Many members of the Forestry Department were closely involved in project work on the biology of the iguana on Saint Lucia, including especially Alwin Dornelly, Timotheus Jean Baptiste, Brian James, Lyndon John, Stephen Lesmond, Michael Bobb, Adams Toussaint, Michael Andrew, Feria Narcisse-Gaston, Rebecca Rock, George Antoine, Mary James and Richard Regis. Martin Satney and Dunley Auguste provided much appreciated support from the Ministry of Agriculture. Special thanks are due to Donald Anthony (Saint Lucia Forestry Department), John Hartley and John Fa (Durrell Wildlife Conservation Trust) for developing the project work on Saint Lucia, and most especially to Anthony ‘Seako’ Johnny for countless hours of assistance with field work and sharing his extensive local knowledge. Many hours of field assistance in Saint Lucia were also contributed by Bradley Abraham, Curtis Mathurin, Neil Oculi, Nazza Gustave, Fendley Estephane, Kissinger Henry and Greg Alexander from Saint Lucia, along with the tireless voluntary efforts of most of the co-authors of Morton *et al.* (2007) plus Jane Huston. Catherine Stephen, Bill Toone, Rich Young, Sarah Seymour, Roger Graveson, Melvin Smith, Chris Pilgrim and most especially Karen Graham all provided much-valued support and insight. The farmers at Sankofa Rainbow Roots Farm and Roots Farm Zimbabwe, along with the owners of Louvet Estate, the Laule family, kindly facilitated access to their lands. This part of our work was funded by the Balcombe Trust.

Pictures and measurements of MCZ specimens were kindly provided by Joseph Martinez. We thank two anonymous reviewers for their remarks on an earlier version of this manuscript.

